# Expression of inhibitory checkpoint ligands by Glioblastoma Multiforme cells and the implications of an enhanced stem cell-like phenotype

**DOI:** 10.1101/2019.12.19.883397

**Authors:** Laverne D Robilliard, Wayne Joseph, Graeme Finlay, Catherine E Angel, E Scott Graham

## Abstract

Glioblastoma Multiforme is a highly aggressive brain malignancy commonly refractory to classical and novel chemo-, radio- and immuno-therapies, with median survival times of ~15 months following diagnosis. Poor immunological responses exemplified by the down-regulation of T-cell activity, and upregulation of immunosuppressive cells within the tumour micro-environment have limited the effectiveness of immunotherapy in GBM to date. Here we show that GBM cells express a large repertoire of inhibitory checkpoint ligands. Furthermore, GBM cells with an enhanced stem cell-like phenotype exhibit heightened levels of inhibitory checkpoint ligands, compared to non-stem cell-like GBM cells. Understanding how GBM modulates an extensive repertoire of immune checkpoint ligands and the functional consequence on immune evasion are necessary to develop effective immuno-therapeutics.

## Background

Glioblastoma Multiforme (GBM) is classified as a WHO Grade IV astrocytoma that continues to circumvent classical and novel chemo-, radio- and immuno-therapies through extensive intratumoral heterogeneity (1, 2). Like the surrounding central nervous system (CNS) tissue GBM tumours exhibit intrinsic complexity through the presence of interacting microglia, macrophages, astrocytes, oligodendrocytes, neurons, glial and neuronal progenitor cells, pericytes, and endothelial cells (1). In particular, the identification of a subpopulation of cells that share features reminiscent of neural stem cells was first described by Singh et al. (2004), who demonstrated that CD133^+^ cells isolated from human GBM tumours were unique in their ability to self-renew and recapitulate parent tumours in mouse xenograft assays (3). The importance of stem cell-like populations residing within GBM tumour microenvironments has been shown, with cancer stem cell populations demonstrating the propensity to initiate and maintain tumour growth, promote immune evasion, enhance intratumoral angiogenesis, and desensitise GBM to radio- and chemo-therapies (4, 5). Clinically, the presence of GBM cancer stem cells (gCSCs) is associated with progression from low grade to high grade gliomas, in part due to vast cancer stem cell interactomes (6). Specifically, GBM tumours actively interact with immune cell populations, potentially through multiple immune checkpoint ligand-receptor interactions.

Immune checkpoint molecules are essential cell-surface receptors utilised by immune cells to mediate intercellular communication (Figure 1) (7). Inhibitory checkpoint receptors serve to negatively regulate the development and effector functions of lymphocyte subsets; namely effector T-cells and natural killer cells, with activation of regulatory T –cells (T_reg_) also reported (7, 8). Through the expression of ligands to checkpoint receptors tumour cells effectively suppress immune reactivity (8). In recent years there has been a growing appreciation that a range of these inhibitory checkpoint ligands are expressed throughout the GBM microenvironment (9, 10). Particularly, the expression of programmed death 1 (PD-1; the cognate receptor to PD-L1) in GBM tumours can be as high as 88% (11). Cytotoxic T-lymphocyte-associated protein-4 (CTLA-4) is an additional checkpoint receptor frequently upregulated by T cells within GBM (12). The presence of PD-1 and CTLA-4 is clinically associated with reduced immunological elimination of malignant cells, primarily through either an increase in T-cell anergy or enhanced T_reg_ mediated immunosuppression. FDA approved therapies (originally for melanoma; ipilimumab, pembrolizumab and nivolumab) aimed at disrupting these inhibitory checkpoint signals are in clinical trials for the treatment of GBM (13). In addition to PD-1 and CTLA-4, other immune checkpoints have been described in GBM, with minimal understanding of their functional relevance (14).

**Figure 1.**
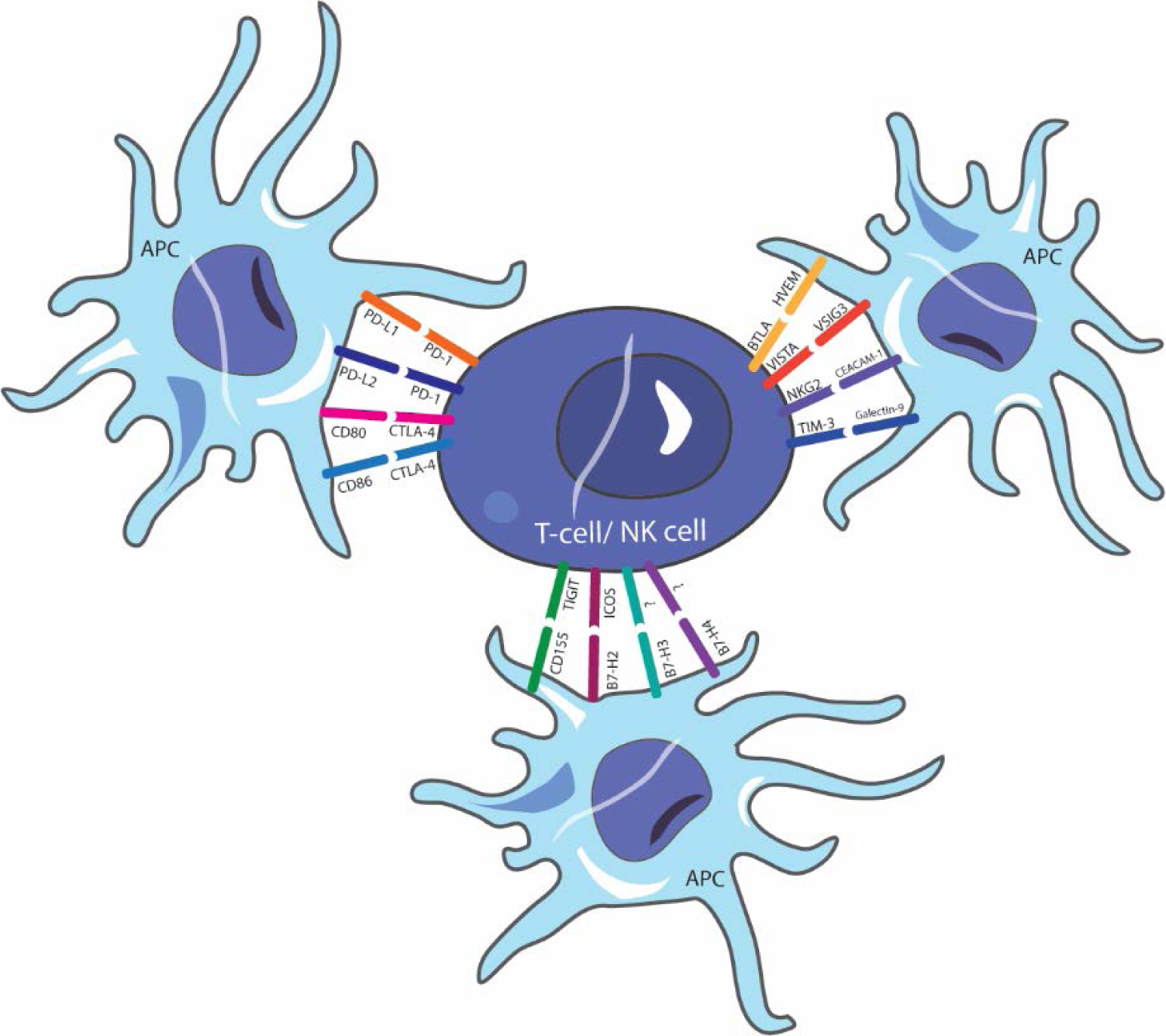
Inhibitory immune checkpoint ligand-receptor interactions. The regulatory suppressive ligands are classically expressed by professional antigen presenting cells (APC) to mediate appropriate immune responses. Cancer cells utilise these mechanisms by aberrantly expressing inhibitory checkpoint ligands as a means of down-regulating anti-tumour immune mechanisms.

An important consideration in the investigation of immune-suppression is the expression and modulation of checkpoint ligands by glioma cancer stem cells. Emerging studies have linked increased PD-L1 expression to CD44^+^ and CD133^+^ CSC populations in non-CNS tumours (14). Wu et al. (2017) demonstrated that both MCF-7 breast cancer and HCT-116 colon cancer cultures enriched for cancer stem cells as defined by CD44^high^CD24^low^ phenotypes exhibit significantly higher percentages of PD-L1 positive cells (15). While the functional implications of elevated checkpoint ligand expression by CSC populations is still unclear, the findings reinforce the multi-faceted role CSCs may play in solid tumour immuno-modulation.

The recent FDA approvals of anti-CTLA-4 and anti-PD-1 immunotherapies have propelled the benefits of targeting immune checkpoints into the spotlight. However, as multiple checkpoint ligands have been demonstrated to play complementary roles in the inhibition of T-cell activity, monotherapies as a means of countering checkpoint mediated immune suppression may not provide the greatest anti-tumour effects (16). For example tumours are able to escape anti-CTLA-4 monotherapy via upregulation of PD-1/PD-L1 interactions (17). It is apparent that inhibitory checkpoint receptor signalling function in concert to mediate inappropriate immune down-regulation. Thus, understanding the extent of checkpoint ligand expression within GBM tumours is necessary to develop functional, long-lasting therapeutics. In this study, we investigate the expression of an extensive range of suppressive checkpoint ligands by two primary New Zealand glioblastoma cell lines. Additionally, we highlight the enhanced expression of checkpoint ligands by stem cell-like enriched populations and recognise the implications of cancer stem cells for future immunotherapeutic interventions.

## Methods

### Cell culture

#### Primary New Zealand Glioblastoma cell lines

NZB11 and NZB19 primary cell lines were provided in collaboration with the Auckland Cancer Society Research Centre. The cells were acquired at a low passage and routinely cultured at 37 °C in 5% O_2_, 5% CO_2_. Cells were cultured as adherent monolayers on uncoated 75 cm^2^ culture flasks until 80-90% confluent. For experimental conditions requiring fetal bovine serum (FBS), cells were cultured in α Minimal Essential Medium (MEM) (ThermoFisher) supplemented with 5% FBS (Moregate) and 1x insulin-transferrin-selenium (ITS) (Sigma) (herein referred to as serum-cultures).

#### Adherent GBM Cancer Stem Cell-Like Cells (gCSC)

Adapted from established glioma stem cell protocols (18), adherent gCSCs were expanded for experimental use and routinely cultured at 37 °C in 5% O_2_, 5% CO_2_. NZB11 and NZB19 primary cell lines at low passages were transferred into 25 cm^2^ culture flasks coated with 10 µg/mL laminin (ThermoFisher). gCSC culture medium consisted of Dulbecco’s Modified Eagle Medium/F12 (DMEM/F12) (ThermoFisher) supplemented with 0.5x B-27 minus vitamin A (ThermoFisher), 0.5x N2 (ThermoFisher), 20 ng/mL bFGF (Peprotech) and 20ng/mL EGF (Novus Biologicals) (herein referred to as gCSC cultures). Half-volume medium changes were carried out every 3 days, for a minimum of 21 days prior to experimental use.

#### Glioma-sphere Formation

For the formation of gCSC glioma-spheres, gCSC cultures were removed from laminin coated flasks and cultured in DMEM/F12 supplemented with 0.5x B-27 minus vitamin A, 0.5x N2, 20 ng/mL bFGF and 20ng/mL EGF. Spheres were routinely cultured at 37 °C in 5 % O_2_, 5% CO_2_.

#### NT2-Astrocytes (NT2A)

NT2A cells were generated as described previously (19) and were cultured on uncoated 25 cm^2^ culture flasks in DMEM/F12 at 37 °C in 20% O_2_, 5% CO_2_.

#### Imaging

For phase imaging of each respective cell culture, cells were imaged at equivalent confluences on EVOS FL auto imaging system (ThermoFisher) at either 10x or 20x magnification.

### Limited Dilution Assay

As per sphere forming conditions, gCSC NZB11 and NZB19 primary cell lines were grown in gCSC culture medium in the absence of laminin coating. To determine gCSC self-renewal potential, primary GBM glioma-spheres were treated with Accutase and triturated until all cells were in single cell suspension. Single cells derived from primary spheres were then seeded into 96 well plates at a cell seeding density of 5 cells per µL. After 2 weeks the number of spheres greater than 80 µm in diameter were counted. Sphere forming efficiency (SFE) was determined by calculating *(No. spheres < 80 µm / No. cells seeded)*100*.

### Flow Cytometry

The expression of surface stem cell-associated molecules and checkpoint ligands were determined by flow cytometry. Primary cell lines were cultured in 75 cm^2^ culture flasks under respective culture conditions until 80-90% confluent. Medium was removed and cells were washed with 1x PBS (ThermoFisher). Following removal of PBS, cells were treated with Accutase (Sigma) for 5 min at room temperature until all cells were in single cell suspension. Accutase was diluted 1:2 with warm medium and suspended cells were centrifuged for 5 min at 300 × g. The supernatant was discarded, and the cells were re-suspended in medium. For flow cytometry preparation, 100,000 cells were added to round-bottom polystyrene tubes for antibody incubations. Cells were incubated at a 1:20 dilution (5 µL into 100 µL cell suspension) for 15 min at 4°C with conjugated primary antibody (Table 1). 7-AAD (BioLegend) was used for live/dead discrimination. Following incubations, each tube was washed once with 2 mL FACS buffer (PBS containing 1% FBS) and centrifuged for 5 min at 300 × g at 4 °C. Supernatant was decanted and cells were vortexed vigorously immediately prior to flow cytometry analysis. Cells were analysed using a BD Accuri C6 Flow Cytometer. Data was processed using FlowJo v.7.6.5 software. Dot plots were created in GraphPad Prism v.7.

**Table 1.**
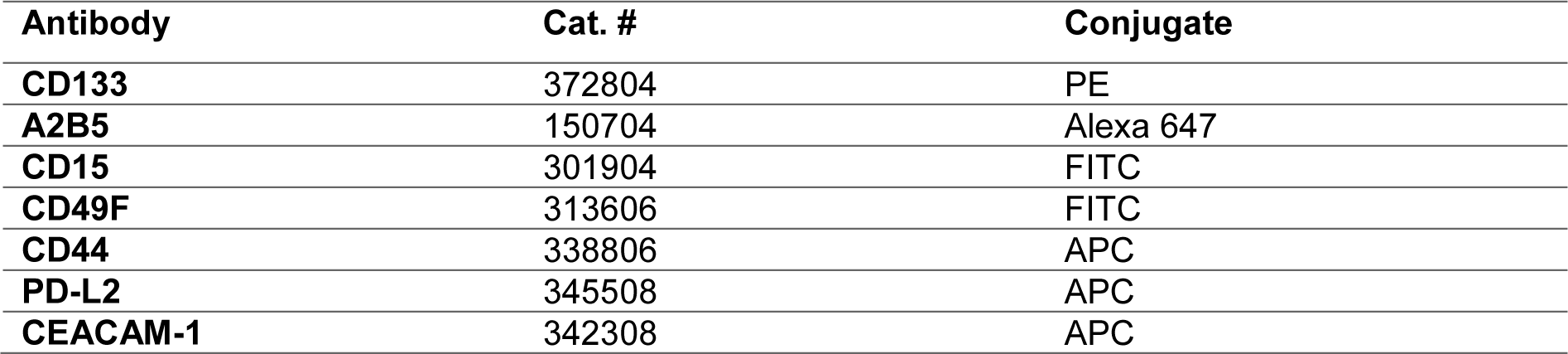
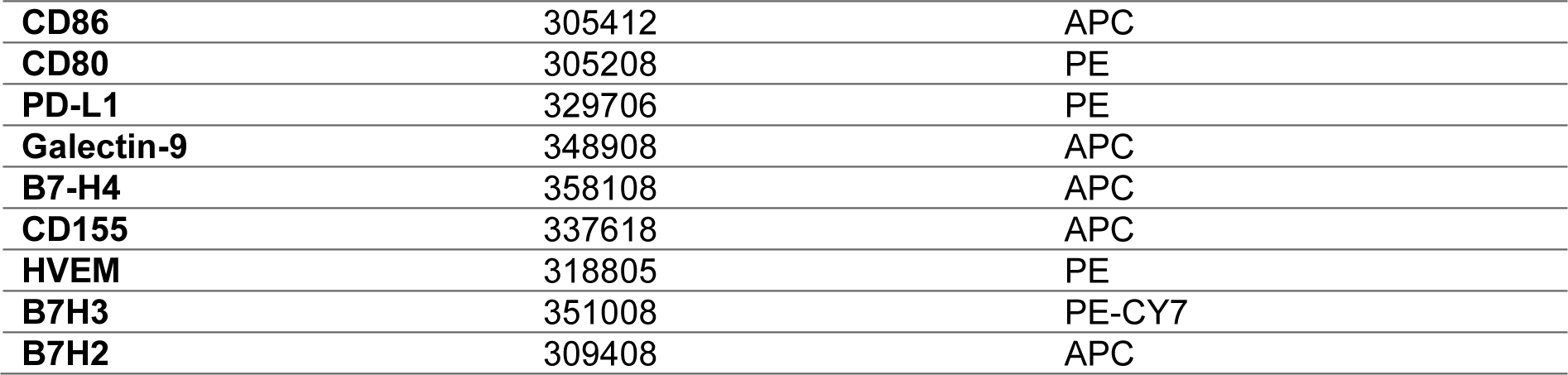
Conjugated primary antibodies used for flow cytometry

| Antibody | Cat. # | Conjugate |
| --- | --- | --- |
| CD133 | 372804 | PE |
| A2B5 | 150704 | Alexa 647 |
| CD15 | 301904 | FITC |
| CD49F | 313606 | FITC |
| CD44 | 338806 | APC |
| PD-L2 | 345508 | APC |
| CEACAM-1 | 342308 | APC |
| **CD86** | 305412 | APC |
| **CD80** | 305208 | PE |
| **PD-L1** | 329706 | PE |
| **Galectin-9** | 348908 | APC |
| **B7-H4** | 358108 | APC |
| **CD155** | 337618 | APC |
| **HVEM** | 318805 | PE |
| **B7H3** | 351008 | PE-CY7 |
| **B7H2** | 309408 | APC |

### Immunocytochemistry

Cells cultured in serum and gCSC medium were seeded at 5000 cells/0.33cm^2^ into either uncoated or laminin coated 96 well plates, respectively. Cells were allowed to settle and continue to proliferate for a further 48 hrs and then fixed. Cells were fixed in either 4% paraformaldehyde (PFA) or 95% methanol, 5% acetic acid for 10 min and then washed once with 1x PBS. Cells fixed in PFA were permeabilised for a further 10 min in 0.1% PBS-Triton X-100 (PBS-T). Cells were washed and stored in PBS. For immunocytochemical analysis, PBS was aspirated, and cells were blocked in 1% Bovine Serum Albumin (BSA) for 45 min and washed thrice for 10 min in 0.1% PBS-T. Primary antibodies were diluted, as per table 2, in 1% BSA and incubated with cells for 1 h at room temperature on a rocker. After primary antibody incubations, cells were washed as previously described and incubated with either 1:400 goat anti-mouse 488 conjugated secondary (Cat. #A11001) or 1:400 goat anti-rabbit 594 conjugated secondary (Cat. #A11005) made up in 1% BSA for 1 hr at room temperature. Cells were counterstained with 1:10,000 Hoechst 33342 (ThermoFisher). Cells were then washed as previously described and stored in PBS. For imaging, EVOS FL auto imaging system (ThermoFisher) was used and ImageJ was used to process images.

**Table 2.** Primary antibodies used for immunocytochemistry

| **Antibody** | **Cat. #** | **Dilution** | **Fixation** | **Host** |
| --- | --- | --- | --- | --- |
| **Nestin** | sc-23927 | 1:100 | PFA | Mouse |
| **A2B5** | 150704 | 1:400 | PFA | Mouse |
| **Vimentin** | ab20346 | 1:1000 | Methanol | Mouse |
| **βIII Tubulin** | T8660 | 1:400 | PFA | Mouse |
| **GFAP** | Z0334 | 1:2000 | Methanol | Rabbit |
| **NeuN** | MAB377 | 1:100 | PFA | Mouse |
| **CD44** | ab189524 | 1:400 | Methanol | Rabbit |

### Confocal Microscopy

Glioma-spheres were added to 16-well chamber slides (ThermoFisher) coated with 1:50 Matrigel. Spheres were cultured for 1 hr until they adhered loosely. Spheres were fixed in either 4% PFA or 95% methanol, 5% acetic acid for 10 min and then washed once with 1x PBS. Spheres fixed in PFA were permeabilised for a further 10 min in 0.1% PBS-Triton X-100 (PBS-T). Spheres were washed and stored in PBS. For immunocytochemical analysis, PBS was aspirated and spheres were blocked in 1% Bovine Serum Albumin (BSA) for 45 min and washed thrice for 10 min in 0.1% PBS-T. CD44, GFAP, vimentin, nestin and BIII Tubulin primary antibodies were diluted, as indicated in Table 2, in 1% BSA and incubated with spheres for 1 h at room temperature on a rocker. After primary antibody incubations, spheres were washed as previously described and incubated with either 1:400 goat anti-mouse 488 conjugated secondary (Cat. #A11001) or 1:400 goat anti-rabbit 594 conjugated secondary (Cat. #A11005) made up in 1% BSA for 1 hr at room temperature. Spheres were counterstained with 1:10,000 Hoechst 33342 (ThermoFisher). Spheres were then washed as previously described and stored in PBS. For imaging, an Olympus FV1000 confocal microscope was used and ImageJ was used to process images.

### *In silico* analysis of GBM checkpoint ligand expression

Search strategy used SCOPUS to search for key terms related to GBM, PD-L1, PD-L2, CD80, CD86, Galectin-9, CEACAM-1, CD155, B7-H2, B7-H3, B7-H4 and HVEM on and before 25 March 2019 (Table 3). Keywords were all inclusive. All studies that included one of the search terms within the title, abstract or keywords were included. Duplicates were removed using EndNote Software X8. Only studies that included i) human analysis, ii) mRNA and/or protein detection, and/or iii) direct tumour cell expression of ligands were included.

**Table 3.** *In* silico analysis search terms

| Keyword | Search Term |
| --- | --- |
| **GBM** | "Glioma" OR "GBM" OR "Glioblastoma*" OR "Grade IV astrocytoma*" OR "Glioblastoma Multiforme" OR "Glial tumour" |
| **CD80** | "CD80" OR "B7-1" OR "BB1" OR "CD28LG" OR "CD28LG1" OR "LAB7" |
| **CD86** | "CD86" OR "B7-2" OR "B7.2" OR "B70" OR "CD28LG2" OR "LAB72" |
| **PD-L1** | "CD274" OR "B7-H" OR "B7H1" OR "PD-L1" OR "PDCD1L1" OR "PDL1" OR "Programmed cell death 1 ligand 1" |
| **PD-L2** | "PDCD1LG2" OR "B7DC" OR "Btdc" OR "CD273" OR "PD-L2" OR "PDL2" OR "programmed cell death 1 ligand 2" OR "PDCD1" OR "CD279" |
| **CEACAM-1** | "CEACAM1" OR "BPG" OR "CD66" OR "Carcinoembryonic antigen related cell adhesion molecule 1" OR "NKG2" |
| **CD155** | "PVR" OR "CD155" OR "HVED" OR "NECL5" OR "PVS" OR "TAGE4" OR "poliovirus receptor" |
| **Galectin-9** | "LGALS9" OR "HUAT" OR "LGALS9A" OR "Galectin 9" OR "Galectin-9" |
| **B7-H3** | "CD276" OR "4IG-B7-H3" OR "B7-H3" OR "B7H3" OR "B7RP-2" OR "CD276 molecule" |
| **B7-H2** | "B7-H2" OR "ICOSLG" OR "B7H2" OR "B7HRP-1" OR "B7HRP1" OR "CD275" OR "ICOS-L" OR "LICOS" OR "ICOS LIGAND" |
| **H7-H4** | "VTCN1" OR "B7-H4" OR "B7H4" OR "B7S1" OR "B7X" OR "PRO1291" OR "VCTN1" OR "V-set domain containing T cell activation inhibitor 1" |
| **HVEM** | "TNFSF14" OR "ATAR" OR "CD270" OR "HVEA" OR "HVEM" OR "LIGHTR" OR "TR2" OR "TUMOUR NECROSIS FACTOR RECEPTOR SUPERFAMILY MEMBER 14" OR "TNF RECEPTOR SUPERFAMILY MEMBER 14" OR "BTLA" OR "CD272" OR "B and T lymphocyte associated" |

### Statistical Analysis

Each experiment was repeated three times with representative data shown where applicable. For flow cytometry data, median fluorescent intensities across three independent repeats are presented with students T-tests used for statistical comparisons. GraphPad v.7 was used to generate statistical tests. P = 0.05 (*), 0.01 (**), 0.001 (***), 0.0001(****).

## Results

### Primary New Zealand GBM cell lines readily form adherent cells and free floating glioma-spheres in gCSC culture

To establish GBM cancer stem cell-like cells (gCSCs) *in vitro*, the absence of serum in combination with key mitogens such as EGF and bFGF is required (18). To assess the ability of primary New Zealand GBM cell lines to be grown as gCSCs, NZB11 and NZB19 primary cell lines were cultured and characterised in both serum and serum-free medium. Primary cell lines were initially established from reported glioblastoma resections and cultured in medium supplemented with 5% FBS. These cultures readily expand as adherent, heterogeneous populations in the absence of extracellular matrix. The cells form a mixture of large, flat circular cells reminiscent of non-reactive astrocytes cultured *in vitro* and a smaller population of elongated, spindle-like cells (Figure 2) (18, 20). The heterogeneity of the serum-cultured cells is exemplified by the large range in forward scatter indicated by flow cytometry analysis, demonstrating that the cells exist as an extensive range of cell sizes (Figure S1). The presence of a heterogeneous population likely reflects *in vivo* conditions, whereby the bulk of the primary tumour will contain cells of various phenotypes. However prolonged exposure to serum will undoubtedly promote cellular differentiation and escape from parental phenotypes. Serum-cultured cells exhibit stable growth rates over extended passages and consistently had doubling times of ~1 week.

**Figure 2.**
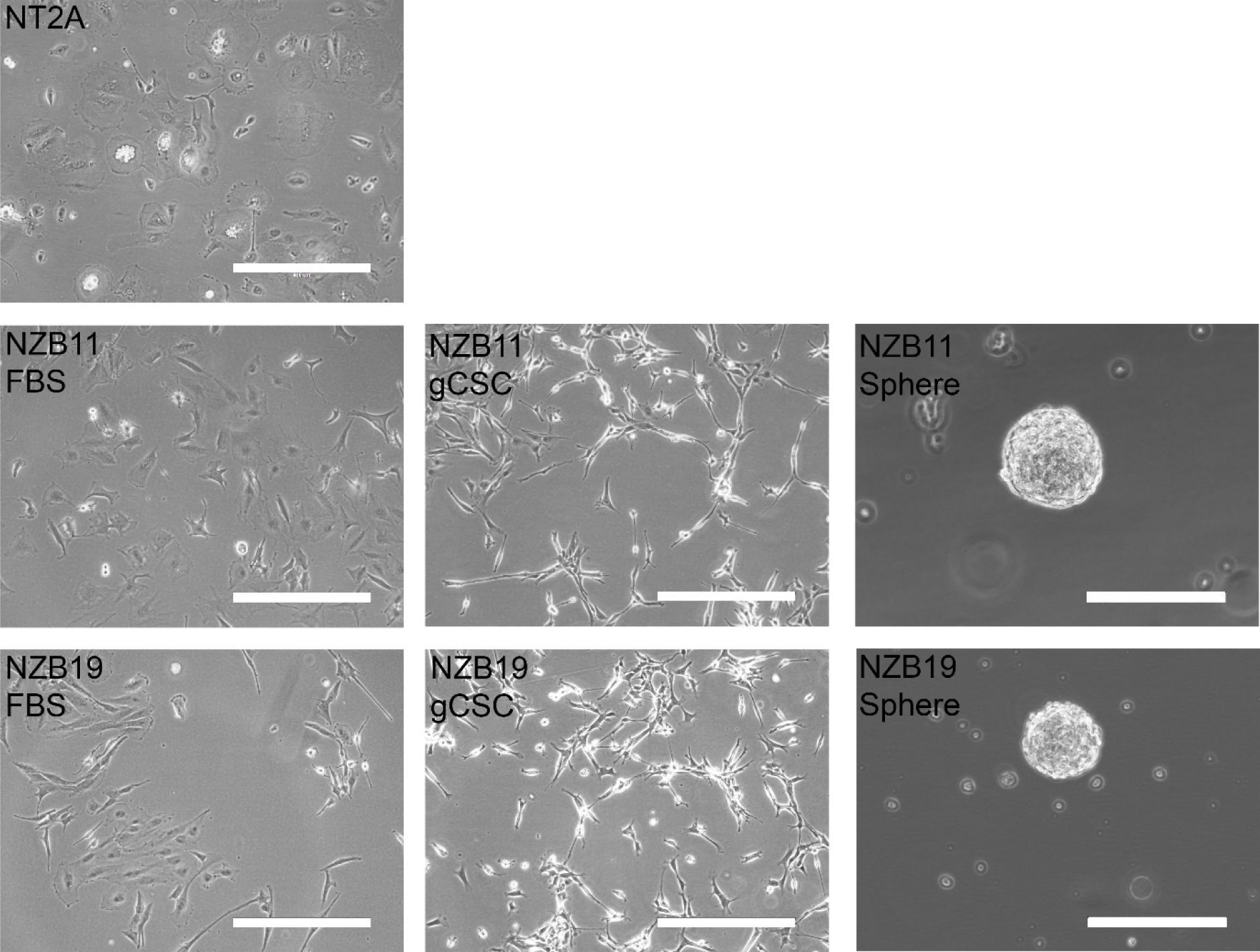
Morphological comparison of New Zealand serum-cultured GBM, g CSC and glioma-sphere cultures. Phase contrast images showing morphological distinctions between NT2 A, NZB11 and NZB19 cell lines in serum-cultured conditions, as adherent gCSC cultures and glioma-spheres. Images were acquired at 10x and 20 × magnification. Adherent culture scale bar = 400 µ m. Glioma-sphere scale bar = 200 µm.

To ascertain whether primary New Zealand GBM cell lines established in serum-containing medium retain stem cell-like potential, at passage ~10, cells were removed from serum and cultured in established gCSC media on 10 µg/mL laminin for a period of 21 days prior to characterisation. Primary cell lines rapidly formed homogenous, adherent cultures that were analogous to the long, spindle-like cells present in the serum-cultured cells (Figure 2). These observations are routinely observed across patient derived primary GBM cells and commercial GBM primary cell lines (18, 21). The cells retain this morphology for subsequent passages over months of culture. The cells showed a definite left-ward shift in their forward scatter, with greater proportions of smaller cells compared to their serum-cultured counterparts (Figure S1).

The current gold standard for the identification and expansion of gCSCs from GBM resections is the ability of primary cell lines to form free-floating glioma-spheres, reflective of neural stem sphere formation. To determine the sphere forming properties of gCSC cultures, cells were cultured in gCSC media in the absence of any extracellular matrix. The cells immediately failed to adhere, instead remained in single cell suspension. ~2 days post transfer into laminin free flasks, ~20 % of the single cells began to replicate and form cell aggregates that remained free-floating. After 2 weeks of sphere culture, each glioma-sphere remained viable and continued to expand until spheres were consistently ~80 µm in diameter. At 4 weeks post adherent culture, the spheres no longer proliferated at ~150 – 200 µm in diameter (Figure 2). Any larger, the spheres rapidly settled on to the plastic and began to spread out and adhere loosely via a small number of migrating cells. NZB11 and NZB19 glioma-spheres retained sphere forming abilities over multiple generations with sphere forming efficiencies of 11.2 % and 15.3 %, respectively (Figure S2).

### GBM cancer stem cell-like cells exhibit increased stem cell associated phenotypes compared to serum-exposed cells

To validate the hypothesis that the gCSC adherent cultures promote the development of cancer stem cells, both primary cell lines were assessed using flow cytometry and immunocytochemistry for the expression of key cancer stem cell associated markers and neural lineage differentiation markers.

Both primary cell lines were analysed for their expression of CD133, A2B5, CD44 and CD49F by flow cytometry (Figure 3). All markers were detected at significant in the serum-supplemented medium. When grown as gCSCs, the expression of A2B5 significantly increased in both primary cell lines. While gCSC cultures did not express CD133 and CD49F substantially above serum-cultured cells, there is a definite increase in the overall stem cell-like phenotype of both primary cell lines grown as gCSCs (Figure 3, B). It is important to note that gCSC A2B5 expression was not only elevated, but also showed broader expression profiles according to flow cytometry histograms (Figure 3, A).

**Figure 3.**
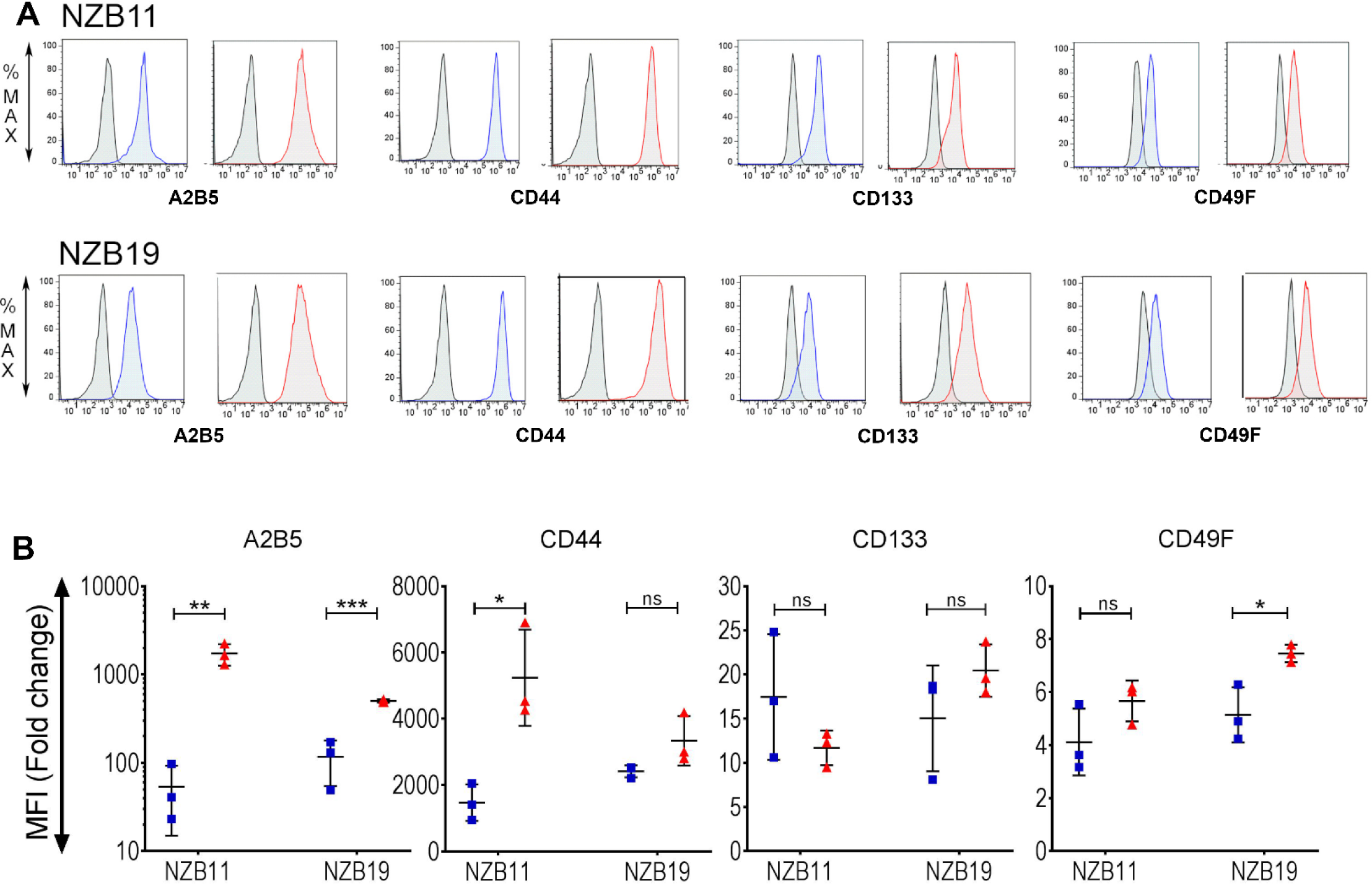
Flow cytometry analysis of cell surface cancer stem cell associated markers. A) Representative histograms of NZB11 and NZB19 cancer stem cell associated marker expression. Blue histograms represent serum-cultured cells, red histograms represent gCSC cultures. Grey histograms represent auto-fluorescence. Shown is one representative experiment of three independent repeats. B) Median fluorescent intensity fold change from auto-fluorescence for each cancer stem cell associated marker. Side by side comparisons of NZB11 and NZB19 serum-cultured vs. gCSC cultured cells. Shown are three independent repeats. Paired students T-test analysis was carried out. P = 0.05 (*), 0.01 (**), 0.001 (***), 0.0001(****).

Next, comparisons between the expression of key stem cell associated and differentiation markers were made by immunocytochemical analysis (Figure 4). Each primary cell line, grown in serum or as gCSCs cultures, were imaged for their expression of nestin, A2B5, vimentin, BIII Tubulin, GFAP and NeuN. All markers assessed were expressed by both primary cell lines (Figure 4). However, gCSC cultures differed from serum-cultured cells in their expression of A2B5. NZB11 and NZB19 gCSC cultures showed increased intensities for A2B5, in agreement with flow cytometry findings (Figure 4, A). Both primary cell lines similarly demonstrated downregulation of BIII Tubulin and NeuN when cultured as gCSCs (Figure 4, B).

**Figure 4.**
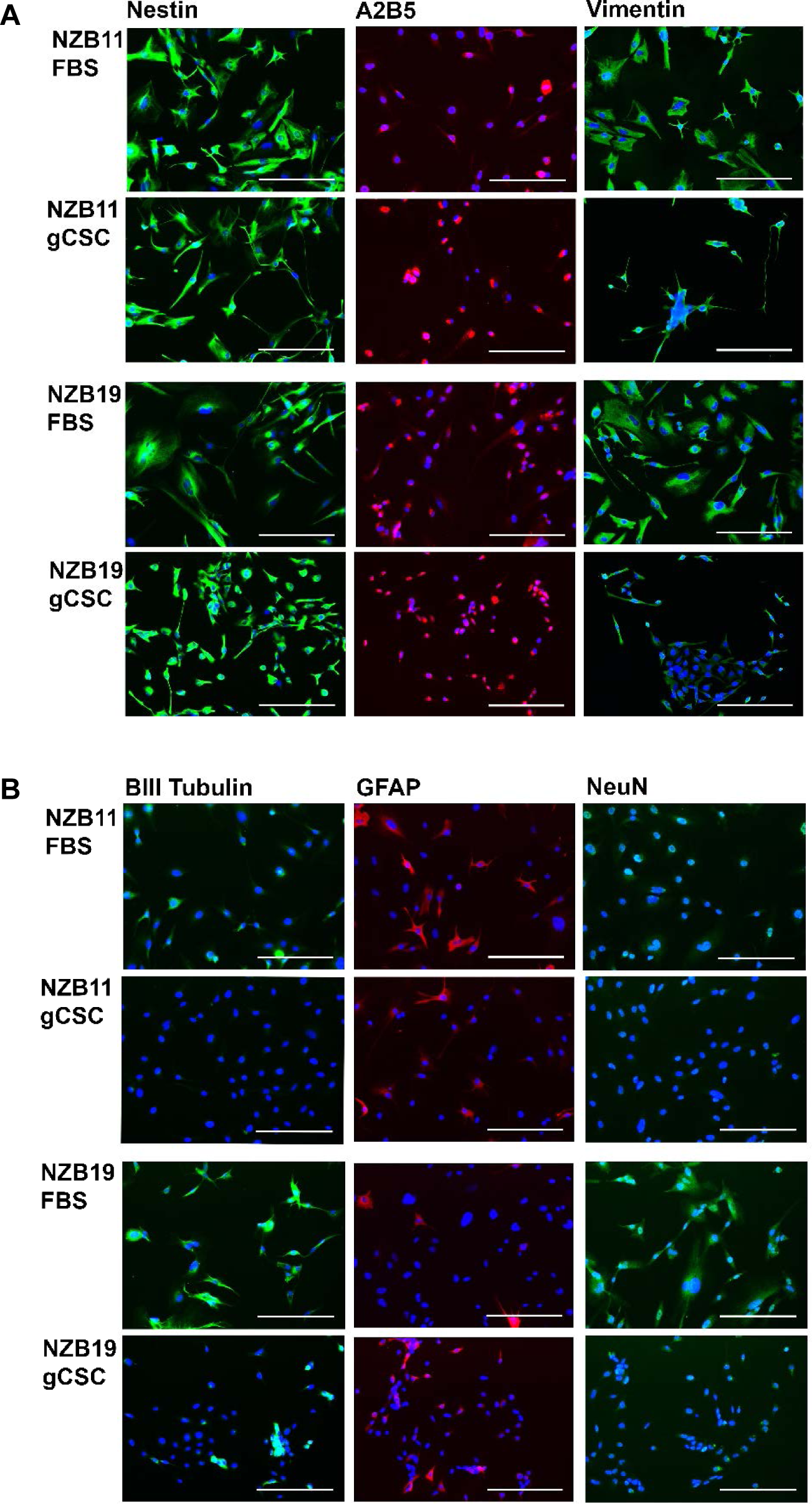
Surface stem cell associated marker and neural lineage differentiation marker expression by GBM cell lines. A) Nestin, A2B5 and Vimentin immuno-staining of NZB11 and NZB19 serum-cultured cells and gCSC cultures. B) BIII Tubulin, GFAP and NeuN immuno-staining of NZB11 and NZB19 serum-cultured cells and gCSC cultures. Cells were seed at 5000 cells/0.33cm^2^ and, fixed 48 hrs post seeding. Images were acquired at x20 magnification. Scale bar = 200 µm. Shown is one representative experiment of three independent repeats.

Furthermore, the expression of nestin, GFAP, CD44, vimentin and BIII Tubulin by GBM derived glioma-spheres was determined (Figure 5). Z-stack analyses revealed that CD44 is expressed throughout glioma-spheres, whereas GFAP, nestin, vimentin and BIII Tubulin are localised to sphere peripheries (Figure S3-S6). The expression profiles between spheres are consistent between both primary cell lines, however the varied expression within each sphere indicate that while a homogenous population of gCSCs forms spheres, the spheres themselves are inherently heterogeneous.

**Figure 5.**
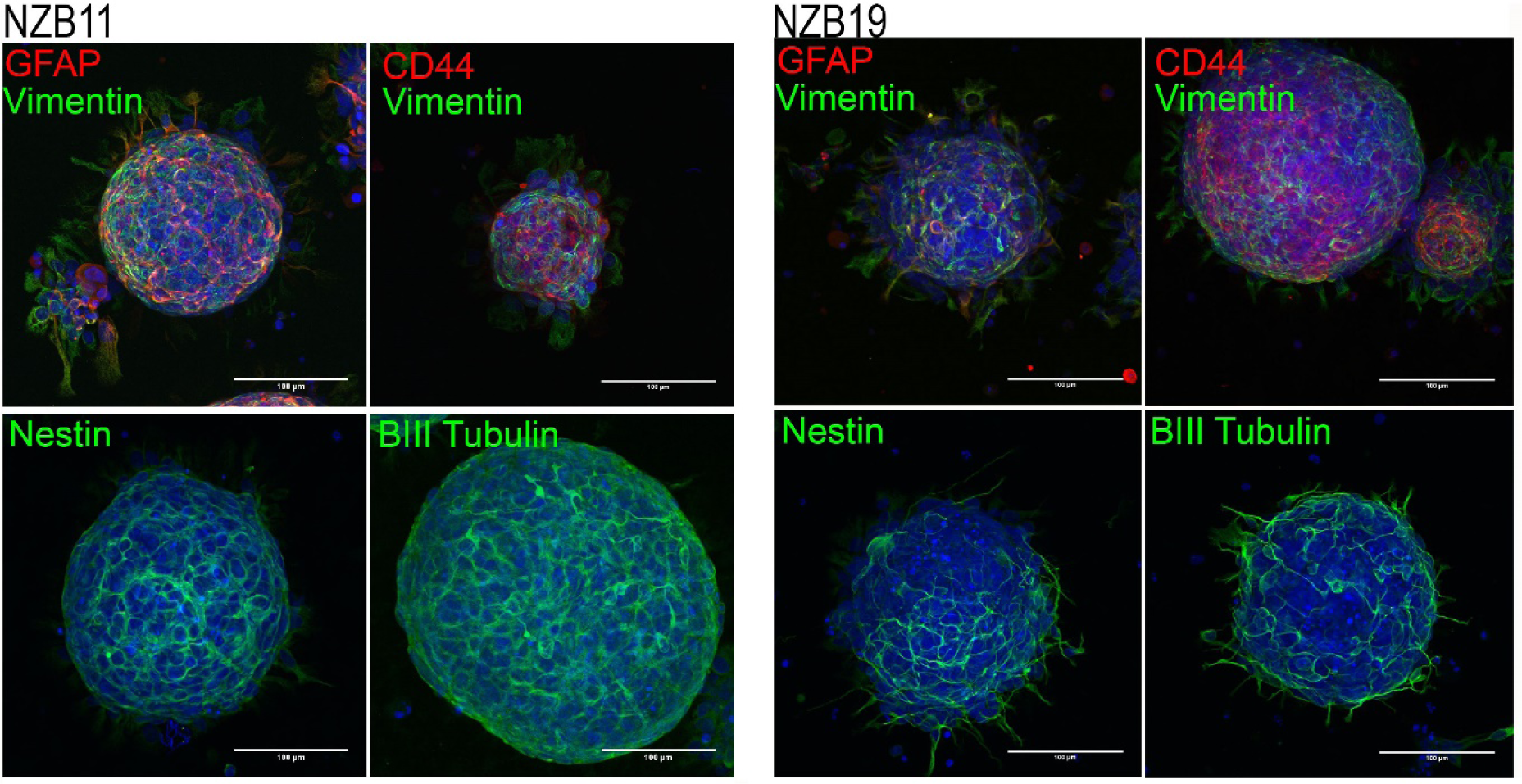
Glioma-sphere expression and localisation of stem cell associated markers. NZB11 and NZB19 gCSC derived glioma-sphere expression of GFAP, vimentin, CD44, nestin and BIII Tubulin. Max intensity Z-project stacks showing the expression of stem-cell associated markers through each glioma-sphere. Images were acquired at x40 magnification. Scale bar = 100 µm.

### Primary New Zealand GBM cells express a wide range of checkpoint ligands that are present at a greater level in gCSCs

The expression of immune inhibitory checkpoint ligands within tumour micro-environments is of immense clinical importance. The expression, however, of these molecules by GBM cancer stem cells is largely unknown. GBM tumours are known to express checkpoint ligands, with expression levels (of PD-L1) often correlated with worsening grades (22). This study is the first to explore the expression of an extensive selection of inhibitory checkpoint ligands on GBM tumour cells and on their cancer stem-cell like counterparts. Figure 6 emphasises the importance of investigating the expression profiles of these molecules as all ligands show rightward shifts in the median fluorescence intensity. Moreover, the broad histogram expression profiles for multiple ligands investigated here is in line with reports that GBM expression of immune modulating molecules are frequently heterogeneous in both gCSC and serum-cultured cells (23). Interestingly, CD80 expression in serum-cultured cells by both primary GBM lines consistently shows skewed rightward expression while gCSC cultures are likely expressing CD80 in two separate populations as indicated by an emerging bimodal profile (Figure 6).

**Figure 6.**
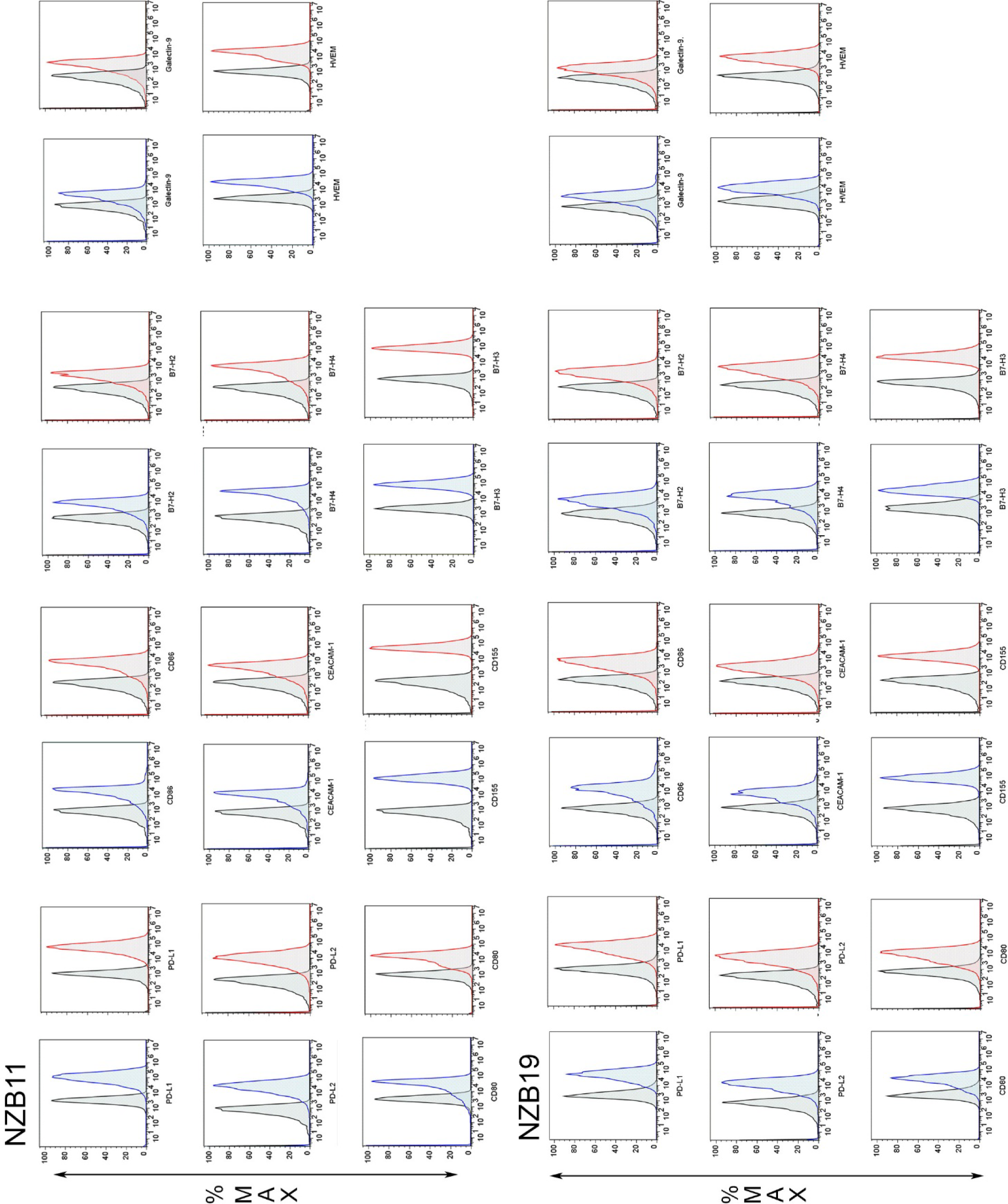
Checkpoint ligand expression in NZB11 and NZB19 primary cell lines. Representative histograms of NZB11 and NZB19 checkpoint ligand expression. Blue histograms represent serum-cultured cells, red histograms represent gCSC cultures. Grey histograms represent auto-fluorescence. Shown is one representative experiment of three independent repeats.

Next, we compared serum-supplemented and gCSC median florescence intensities of the 11 checkpoint ligands against a glial control (NT2A). NZB11 cells express PD-L1 (p=0.0068), PD-L2 (p=0.0002), CD80 (p=0.0083), CD155 (p=0.0046), B7-H2 (p=0.0313), B7-H3 (p=0.0058) and HVEM (p=0.0003) (Figure S8). NZB19’s show expression of PD-L1 (p=0.0277), PD-L2 (p=0.0033), CD80 (p=0.0058), CEACAM-1 (p=0.0057), CD155 (p=0.0030), Galectin-9 (p=0.0097), B7-H3 (p=0.0051) and HVEM (p=0.0264) (Figure S8). The flow cytometry analysis reveals the trend that checkpoint ligands are commonly expressed at higher levels than are seen with non-transformed neural cells, with NZB11’s and NZB19’s positive for 63.6 % and 72.7 % of inhibitory checkpoint ligands, respectively (Figure S8).

As the expression of the aforementioned ligands has been shown to be associated with worsening grade in gliomas and some cancer stem cells in non-CNS tumours, we sought to investigate the expression of these molecules by GBM gCSC cultures. Of the 11 ligands, both the NZB11 and NZB19 gCSCs showed greater levels of surface expression of 3 of the 11 ligands than that of serum-cultured cells (Figure 7). In particular, NZB11 gCSCs expressed surface PD-L1, CD155 and B7-H3 at 23.3, 73.2 and 65.0 fold above that of serum-cultured cells, respectively (Figure 7). In addition, there was a slight increase (not significant) for PD-L2, CD80, Galectin-9, HVEM, and CD86 in the gCSC cultures.

**Figure 7.**
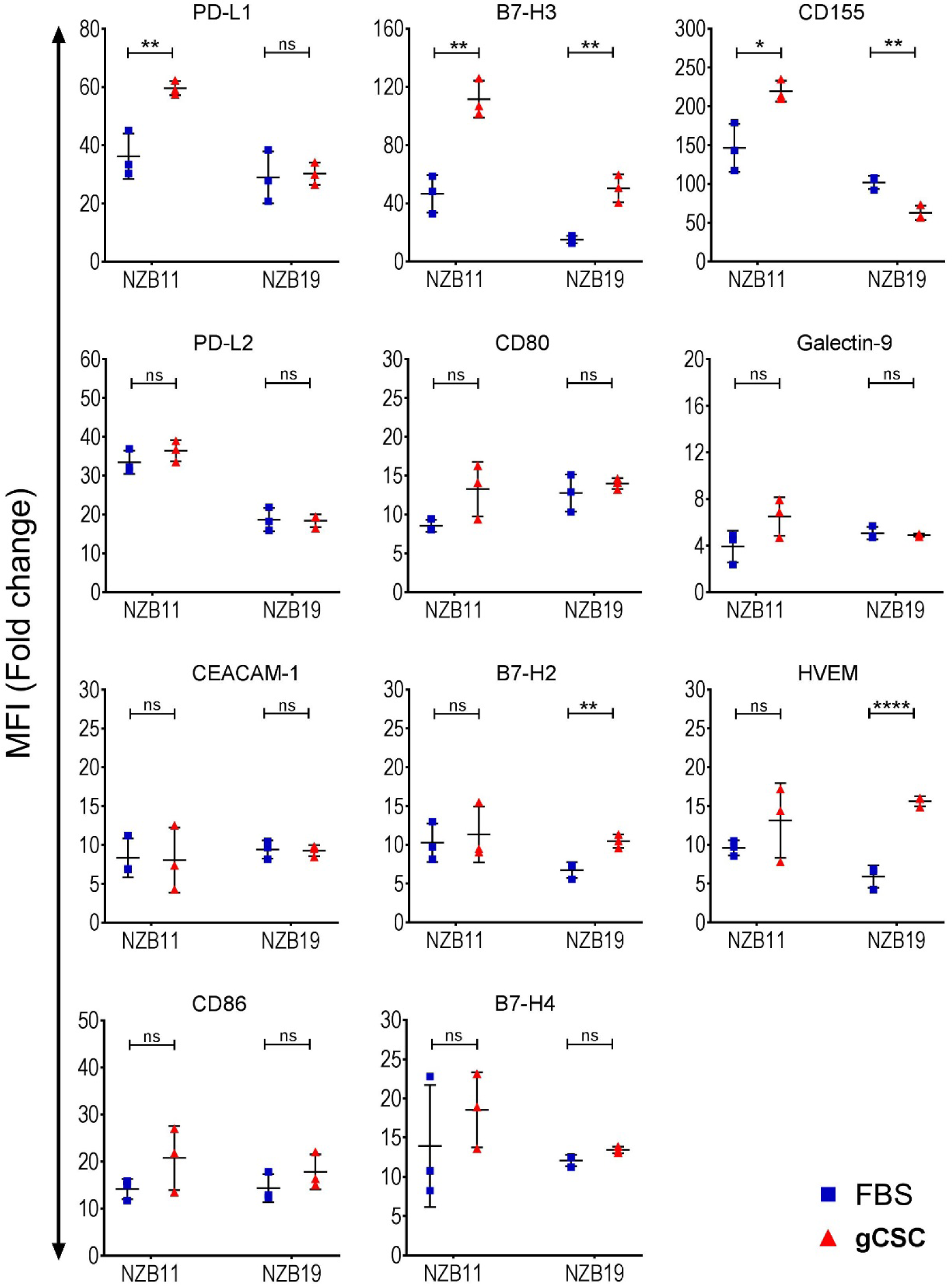
Checkpoint ligand expression analysis across two primary New Zealand GBM cell lines grown as gCSCs. Median fluorescent intensity fold change from auto-fluorescence for each checkpoint ligand. Side by side comparisons of NZB11 and NZB19 serum-cultured cells vs. gCSC cultures. Shown are three independent repeats. Paired students T-test analysis was carried out. P = 0.05 (*), 0.01 (**), 0.001 (***), 0. 0001(****).

Of interest was the observation that the NZB19 gCSC cultures had significantly more B7-H3, B7-H2, and HVEM, but there was no change in PD-L1 or PD-L2. Also contrary to other NZB11 gCSC changes, the surface expression of CD155 was reduced in the gCSC NZB19 cell cultures. (Figure 7). This is very intriguing as not all ligands are elevated, and it is not the same complement of ligands changing across these two distinct lines. Clinically, very few of these molecules are currently being targeted and typically are only targeted by monotherapy. This data reinforces the urgent need to consider checkpoint ligands as a collective group that are differentially regulated and expressed across patient derived glioblastomas.

Figure 8 successfully demonstrates the correlation between increased stem cell phenotypes and elevated expression of inhibitory checkpoint ligands. The role of cancer stem cells in checkpoint immuno-modulation has only very recently been considered outside of non-CNS tumours. Taken together, these results indicate that checkpoint ligands are differentially expressed within glioblastoma in response to increased stem cell-like phenotypes.

**Figure 8.**
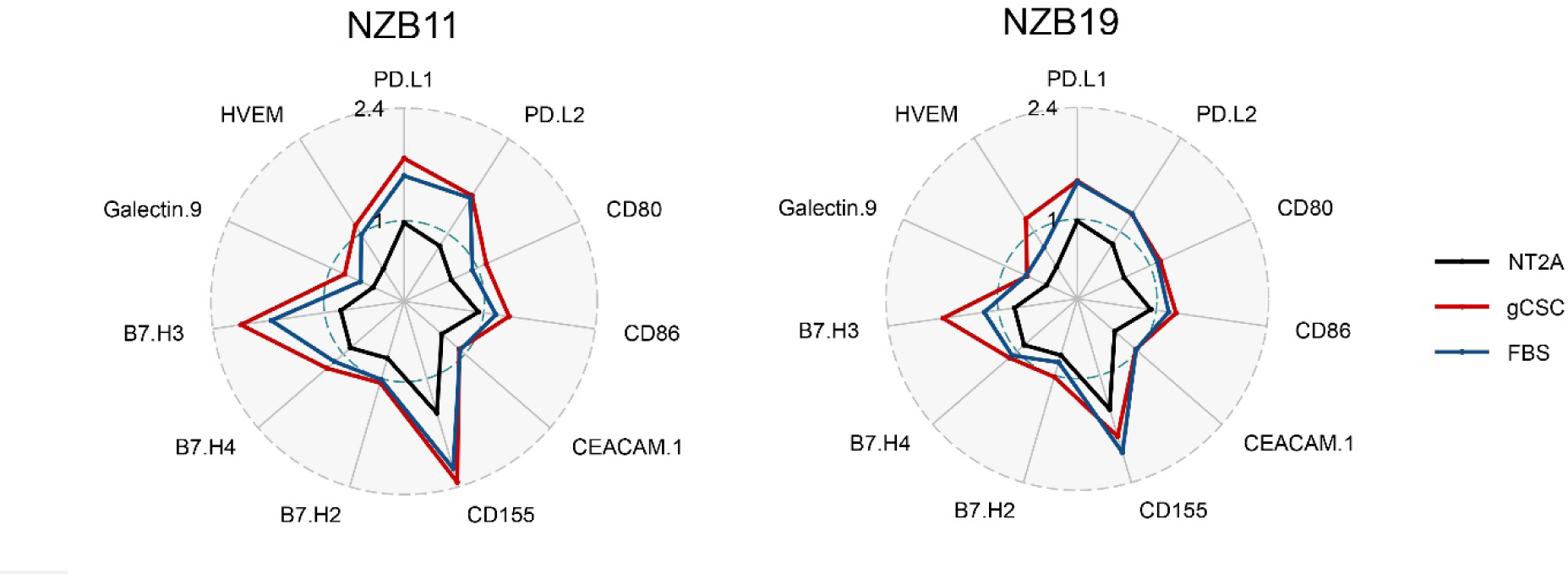
Visualisation of checkpoint ligand expression by New Zealand Glioblastoma cells. Radar plot representation of flow cytometry median fluorescent intensity fold change from auto-fluorescence in NT2A’s, NZB11’s and NZB19’s cultured in serum or as gCSCs (n=3). Scale = Log^10^.

## Discussion

The resounding successes of anti-PD-1 (nivolumab) and anti-CTLA-4 (ipilimumab) in maintaining durable response in melanoma led to the start of a range of clinical trials in GBM (16, 24). Successful treatment of GBM has remained modest at best, with immunotherapies against immune checkpoint ligand-receptor interactions failing to translate into adequate clinical responses. At the forefront of GBM immune checkpoint inhibition was the phase 3, randomised CHECKMATE-143 trial. While the trial failed to induce enhanced overall response rates (8%) in comparison to bevacizumab; an anti-angiogenic (23%), further clinical trials arising from the original concept are underway (25). In particular, trials utilising the combination of nivolumab with either temozolomide, radiation or ipilimumab have commenced, with results pending (26–28). However, the benefits of immune checkpoint inhibition has been limited, primarily due to the commonly low GBM mutational burden and the frequent depletion of lymphocytes within the tumour microenvironment (29, 30). One investigation of a small cohort (n=32) of patients with recurrent glioblastoma aimed to overcome the commonly low immune-response by modifying the immune microenvironment. Through the administration of the anti-PD-1 monoclonal antibody; pembrolizumab in a neoadjuvant fashion, the investigation successfully induced functional activation of tumour infiltrating lymphocytes (TILs), increased tumour-specific T cell clones and caused systemic phenotypic changes in CD4+ T cells to upregulate CD152 and CD127 with concurrent decreases in PD-1 expression. Furthermore, patients showed prolonged overall survival indicating the importance of appropriate T cell activity for effective checkpoint therapies (31). While promising, these studies typically don’t consider the importance of the expression of multiple, likely interacting, immune checkpoint molecules beyond those operating in the PD-1/CTLA-4 axes.

Presented here is, to the best of our knowledge, the first extensive characterisation of surface expression of immune checkpoint ligands with known immune suppressive functions (Figure 6). Surface expression analyses of 11 checkpoint ligands reputed to be immunosuppressive within tumour microenvironments revealed that primary New Zealand GBM cells ubiquitously express these ligands (32). An *in-silico* review of glioblastoma literature revealed that all ligands have been investigated, except for HVEM (Table S1). However, to date investigations have been heavily skewed towards the PD-L1/PD-1 axis, with limited literature on expression profiles of PD-L2, CD80, CD86, B7-H2, B7-H3, B7-H4, CEACAM-1, HVEM and CD155 in human GBM. Collectively, an *in-silico* search revealed that 55% of checkpoint ligand studies in GBM were centred on PD-L1 expression. Therefore, extending our investigation beyond commonly investigated ligands was of importance. Of the ligands we have investigated here, PD-L1, CD155, PD-L2 and B7-H3 surface expression levels are between 8 to 118-fold higher on GBM cells than on astrocytic controls (Figure S8). All 4 ligands are unanimously reported to be expressed in human GBM (Table S1). Therefore, in lieu of the expanding repertoire of checkpoint ligand expression by GBM, the clinical relevance of tumour immune checkpoint interactions needs to be considered heavily. For example, it has been shown that B7-H3 expression by glioma cells corresponds with malignancy grade, likely due to suppression of natural killer cell mediated cell lysis. Further it has been demonstrated by gene silencing of B7-H3 that B7-H3 negative glioma primary cell lines injected into orthotopic models result in greater susceptibility of tumour cells to natural killer cell lysis (33). Similarly, PD-L2 expression across 1357 glioma samples conferred shorter survival times compared to low expressing counterparts. In addition, PD-L2 expression positively correlated with T-regulatory signatures indicating the functional role of PD-L2 in enhancing glioma immunosuppression (34). A similar study of 976 brain glioma samples showed the same functional implications of PD-L1 expression, whereby WHO grade IV gliomas had higher levels of PD-L1 expression compared to grade II and III gliomas. As per PD-L2, PD-L1 expression positively correlated with T cell immunosuppressive signatures (22). Importantly, Wang et al. (2016) also determined that PD-L1 expression correlated with PD-L2 and CD80 expression, further evincing the higher levels of immune suppression present in glioblastoma compared to lower glioma grades (22). One other study has also demonstrated this correlation, whereby high grade gliomas showed high correlations between PD-L2, CD80 and PD-L1 (35). Collectively, this highlights the impact of checkpoint ligand expression on glioma grade and subsequent immune evasion. However, only 18% of human glioma investigations looked at expression levels of multiple ligands, and frequently only in the scope of PD-1/PD-L1/2 and CTLA-4/CD80/86 axes (Table S1). Clinically, expression of multiple ligands poses challenges in targeting GBM through checkpoint inhibition, as the regulation of these ligands in concert is still unclear.

To maximise the potential of immunotherapies, the mechanisms exploited by GBM to promote enhanced checkpoint ligand expression need to be understood. The intrinsic heterogeneity in not only the classification of GBM, but within the tumour microenvironment heavily influences the poor patient survival and ultimate recurrence that is observed (36). In particular, recurrence is likely due to residual GBM subpopulations of stem cells (37). Currently markers reputed for delineating GBM cancer stem cells within the tumour microenvironment include CD133, CD44, A2B5, CD15 and CD49F (38). GBM cancer stem cells can be broadly classified as pro-neural, non-adherent, non-invasive CD133^+^ populations, or as semi-adherent, invasive mesenchymal CD44^+^ populations (37, 39). Flow cytometry analysis revealed that both NZB11 and NZB19 primary cell line derived gCSCs showed increased A2B5 levels. However, NZB11 gCSCs highly expressed CD44, but did not show an increase in CD133 expression (Figure 3). Conversely, NZB19 gCSCs did not show an increase in CD44, but did exhibit elevated CD49F and a non-significant increase in CD133 (Figure 3). These findings exemplify that GBM stem cells display a range of phenotypic characteristics, reiterated by Brown et el. (2015), despite retaining classic cancer stem cell features such as sphere forming ability (Figure 2) (37). To determine the role cancer stem cell populations may play in promoting immunological blockades, our two distinct gCSC lines were screened for the 11 checkpoint ligands previously described (Figure 7). gCSCs are known to impede immunity through multiple modalities, in particular through the expression of inhibitory checkpoint ligands and consequential T-cell anergy (40). Elevated expression by two gCSC lines of 5 of the 11 checkpoint ligands was shown. Importantly, the distinct gCSCs did not show identical expression patterns. The differential expression levels of the checkpoint ligands indicate that phenotypically different gCSCs may utilise discrete repertoires of inhibitory molecules to mediate immune inhibition. B7-H3, the only commonly elevated ligand by NZB11 and NZB19 gCSCs, expression correlates with glioma grade and negatively correlates with T-cell mediated immune response to glioma (41, 42). While B7-H3 expression has not been well characterised on glioma cancer stem cell subsets beyond this study, the elevated expression here suggests to a mechanism by which B7-H3 expression on GBM cancer stem cells could drive worsening prognosis through enhanced immunological inhibition.

## Conclusions

The data here indicates that an increased GBM stem cell-like phenotype results in elevated expression of a range of checkpoint ligands, beyond the extensive repertoire already expressed by serum-cultured cells. The role of checkpoint ligand expression by GBM cancer stem cell subsets ultimately warrants further investigation. Attempting to target these subsets and the mechanisms they use to actively evade immune detection provides a unique therapeutic opportunity in the context of immunotherapies against Glioblastoma Multiforme.

## List of abbreviations

GBM: Glioblastoma Multiforme
CNS: Central nervous system
gCSC: GBM cancer stem cell
FBS: Fetal bovine serum
SFE: Sphere forming efficiency
BSA: bovine serum albumin
PFA: Paraformaldehyde
TIL: tumour infiltrating lymphocyte
CTLA-4: Cytotoxic T-lymphocyte-associated protein-4
PD-1: Programmed death-1
PD-L1: Programmed death ligand-1
PD-L2: Programmed death ligand-2
CEACAM-1: Carcinoembryonic antigen-related cell adhesion molecule-1
HVEM: Herpesvirus entry mediator

## Declarations

### Consent for publication

All authors have approved manuscript for publication.

## Availability of data and materials

All data generated or analysed during this study are included in this published article [and its supplementary information files]

## References

1. Bonavia R, Inda MM, Cavenee WK, Furnari FB. Heterogeneity maintenance in glioblastoma: a social network. Cancer Res. 2011;71(12):4055–60.

2. Shapiro JR, Yung WK, Shapiro WR. Isolation, karyotype, and clonal growth of heterogeneous subpopulations of human malignant gliomas. Cancer Res. 1981;41(6):2349–59.

3. Singh SK, Hawkins C, Clarke ID, Squire JA, Bayani J, Hide T, et al. Identification of human brain tumour initiating cells. Nature. 2004;432(7015):396–401.

4. Gilbert CA, Ross AH. Cancer stem cells: cell culture, markers, and targets for new therapies. J Cell Biochem. 2009;108(5):1031–8.

5. Lathia JD, Mack SC, Mulkearns-Hubert EE, Valentim CL, Rich JN. Cancer stem cells in glioblastoma. Genes Dev. 2015;29(12):1203–17.

6. Silver DJ, Lathia JD. Revealing the glioma cancer stem cell interactome, one niche at a time. J Pathol. 2018;244(3):260–4.

7. Melero I, Berman DM, Aznar MA, Korman AJ, Gracia JLP, Haanen J. Evolving synergistic combinations of targeted immunotherapies to combat cancer. Nature Reviews Cancer. 2015;15:457.

8. Diesendruck Y, Benhar I. Novel immune check point inhibiting antibodies in cancer therapy— Opportunities and challenges. Drug Resistance Updates. 2017;30:39–47.

9. Li J, Liu X, Duan Y, Liu Y, Wang H, Lian S, et al. Combined blockade of T cell immunoglobulin and mucin domain 3 and carcinoembryonic antigen-related cell adhesion molecule 1 results in durable therapeutic efficacy in mice with intracranial gliomas. Medical science monitor: international medical journal of experimental and clinical research. 2017;23:3593.

10. Hung AL, Maxwell R, Theodros D, Belcaid Z, Mathios D, Luksik AS, et al. TIGIT and PD-1 dual checkpoint blockade enhances antitumor immunity and survival in GBM. Oncoimmunology. 2018;7(8):e1466769.

11. Wöhrer A, Brandstetter A, Kiesel B, Marosi C, Zielinski CC, Widhalm G, et al. Programmed death ligand 1 expression and tumor-infiltrating lymphocytes in glioblastoma. Neuro-Oncology. 2014;17(8):1064–75.

12. Fong B, Jin R, Wang X, Safaee M, Lisiero DN, Yang I, et al. Monitoring of Regulatory T Cell Frequencies and Expression of CTLA-4 on T Cells, before and after DC Vaccination, Can Predict Survival in GBM Patients. PLOS ONE. 2012;7(4):e32614.

13. Kim ES, Kim JE, Patel MA, Mangraviti A, Ruzevick J, Lim M. Immune checkpoint modulators: an emerging antiglioma armamentarium. Journal of immunology research. 2016;2016.

14. Zhang D, Tang DG, Rycaj K, editors. Cancer stem cells: Regulation programs, immunological properties and immunotherapy. Seminars in cancer biology; 2018: Elsevier.

15. Wu Y, Chen M, Wu P, Chen C, Xu ZP, Gu W. Increased PD-L1 expression in breast and colon cancer stem cells. Clin Exp Pharmacol Physiol. 2017;44(5):602–4.

16. Wolchok JD, Kluger H, Callahan MK, Postow MA, Rizvi NA, Lesokhin AM, et al. Nivolumab plus ipilimumab in advanced melanoma. New England Journal of Medicine. 2013;369(2):122–33.

17. Shi LZ, Fu T, Guan B, Chen J, Blando JM, Allison JP, et al. Interdependent IL-7 and IFN-γ signalling in T-cell controls tumour eradication by combined α-CTLA-4+ α-PD-1 therapy. Nature communications. 2016;7:12335.

18. Pollard SM, Yoshikawa K, Clarke ID, Danovi D, Stricker S, Russell R, et al. Glioma stem cell lines expanded in adherent culture have tumor-specific phenotypes and are suitable for chemical and genetic screens. Cell Stem Cell. 2009;4(6):568–80.

19. Raos B, Simpson M, Doyle C, Murray A, Graham E, Unsworth C. Patterning of functional human astrocytes onto parylene-C/SiO2 substrates for the study of Ca2+ dynamics in astrocytic networks. Journal of neural engineering. 2018;15(3):036015.

20. Lee J, Kotliarova S, Kotliarov Y, Li A, Su Q, Donin NM, et al. Tumor stem cells derived from glioblastomas cultured in bFGF and EGF more closely mirror the phenotype and genotype of primary tumors than do serum-cultured cell lines. Cancer cell. 2006;9(5):391–403.

21. Al-Mayhani TMF, Ball SL, Zhao J-W, Fawcett J, Ichimura K, Collins PV, et al. An efficient method for derivation and propagation of glioblastoma cell lines that conserves the molecular profile of their original tumours. Journal of neuroscience methods. 2009;176(2):192–9.

22. Wang Z, Zhang C, Liu X, Wang Z, Sun L, Li G, et al. Molecular and clinical characterization of PD-L1 expression at transcriptional level via 976 samples of brain glioma. OncoImmunology. 2016;5(11).

23. Di Tomaso T, Mazzoleni S, Wang E, Sovena G, Clavenna D, Franzin A, et al. Immunobiological characterization of cancer stem cells isolated from glioblastoma patients. Clinical Cancer Research. 2010;16(3):800–13.

24. Wolchok JD, Chiarion-Sileni V, Gonzalez R, Rutkowski P, Grob J-J, Cowey CL, et al. Overall survival with combined nivolumab and ipilimumab in advanced melanoma. New England Journal of Medicine. 2017;377(14):1345–56.

25. Reardon D, Omuro A, Brandes A, Rieger J, Wick A, Sepulveda J, et al. OS10. 3 randomized phase 3 study evaluating the efficacy and safety of nivolumab vs bevacizumab in patients with recurrent glioblastoma: CheckMate 143. Neuro-oncology. 2017;19(suppl_3):iii21–iii.

26. Omuro A, Vlahovic G, Lim M, Sahebjam S, Baehring J, Cloughesy T, et al. Nivolumab with or without ipilimumab in patients with recurrent glioblastoma: results from exploratory phase I cohorts of CheckMate 143. Neuro-oncology. 2017;20(5):674–86.

27. Sampson JH, Omuro AMP, Preusser M, Lim M, Butowski NA, Cloughesy TF, et al. A randomized, phase 3, open-label study of nivolumab versus temozolomide (TMZ) in combination with radiotherapy (RT) in adult patients (pts) with newly diagnosed, O-6-methylguanine DNA methyltransferase (MGMT)-unmethylated glioblastoma (GBM): CheckMate-498. American Society of Clinical Oncology; 2016.

28. Weller M, Vlahovic G, Khasraw M, Brandes A, Zwirtes R, Tatsuoka K, et al. A randomized phase 2, single-blind study of temozolomide (TMZ) and radiotherapy (RT) combined with nivolumab or placebo (PBO) in newly diagnosed adult patients (pts) with tumor O6-methylguanine DNA methyltransferase (MGMT)-methylated glioblastoma (GBM)—CheckMate-548. Annals of Oncology. 2016;27(suppl_6).

29. Thorsson V, Gibbs DL, Brown SD, Wolf D, Bortone DS, Yang T-HO, et al. The immune landscape of cancer. Immunity. 2018;48(4):812–30. e14.

30. Lawrence MS, Stojanov P, Polak P, Kryukov GV, Cibulskis K, Sivachenko A, et al. Mutational heterogeneity in cancer and the search for new cancer-associated genes. Nature. 2013;499(7457):214.

31. Cloughesy TF, Mochizuki AY, Orpilla JR, Hugo W, Lee AH, Davidson TB, et al. Neoadjuvant anti-PD-1 immunotherapy promotes a survival benefit with intratumoral and systemic immune responses in recurrent glioblastoma. Nature Medicine. 2019:1.

32. Martin-Orozco N, Li Y, Wang Y, Liu S, Hwu P, Liu Y-J, et al. Melanoma cells express ICOS ligand to promote the activation and expansion of T-regulatory cells. Cancer research. 2010;70(23):9581–90.

33. Lemke D, Pfenning PN, Sahm F, Klein AC, Kempf T, Warnken U, et al. Costimulatory protein 4IgB7H3 drives the malignant phenotype of glioblastoma by mediating immune escape and invasiveness. Clinical Cancer Research. 2012;18(1):105–17.

34. Wang ZL, Li GZ, Wang QW, Bao ZS, Wang Z, Zhang CB, et al. PD-L2 expression is correlated with the molecular and clinical features of glioma, and acts as an unfavorable prognostic factor. OncoImmunology. 2019;8(2).

35. Guan X, Zhang C, Zhao J, Sun G, Song Q, Jia W. CMTM6 overexpression is associated with molecular and clinical characteristics of malignancy and predicts poor prognosis in gliomas. EBioMedicine. 2018;35:233–43.

36. Verhaak RG, Hoadley KA, Purdom E, Wang V, Qi Y, Wilkerson MD, et al. Integrated genomic analysis identifies clinically relevant subtypes of glioblastoma characterized by abnormalities in PDGFRA, IDH1, EGFR, and NF1. Cancer cell. 2010;17(1):98–110.

37. Brown DV, Daniel PM, Giovanna M, Gogos A, Ng W, Morokoff AP, et al. Coexpression analysis of CD133 and CD44 identifies proneural and mesenchymal subtypes of glioblastoma multiforme. Oncotarget. 2015;6(8):6267.

38. Glaser T, Han I, Wu L, Zeng X. Targeted Nanotechnology in Glioblastoma Multiforme. Frontiers in Pharmacology. 2017;8(166).

39. Mao P, Joshi K, Li J, Kim S-H, Li P, Santana-Santos L, et al. Mesenchymal glioma stem cells are maintained by activated glycolytic metabolism involving aldehyde dehydrogenase 1A3. Proceedings of the National Academy of Sciences. 2013;110(21):8644–9.

40. Silver DJ, Sinyuk M, Vogelbaum MA, Ahluwalia MS, Lathia JD. The intersection of cancer, cancer stem cells, and the immune system: therapeutic opportunities. Neuro-oncology. 2015;18(2):153–9.

41. Baral A, Ye HX, Jiang PC, Yao Y, Mao Y. B7-H3 and B7-H1 expression in cerebral spinal fluid and tumor tissue correlates with the malignancy grade of glioma patients. Oncology Letters. 2014;8(3):1195–201.

42. Zhang C, Zhang Z, Li F, Shen Z, Qiao Y, Li L, et al. Large-scale analysis reveals the specific clinical and immune features of B7-H3 in glioma. OncoImmunology. 2018;7(11).

43. Bloch O, Crane CA, Kaur R, Safaee M, Rutkowski MJ, Parsa AT. Gliomas promote immunosuppression through induction of B7-H1 expression in tumor-associated macrophages. Clinical Cancer Research. 2013;19(12):3165–75.

44. Lee KS, Lee K, Yun S, Moon S, Park Y, Han JH, et al. Prognostic relevance of programmed cell death ligand 1 expression in glioblastoma. Journal of Neuro-Oncology. 2018;136(3):453–61.

45. Liu Y, Carlsson R, Ambjørn M, Hasan M, Badn W, Darabi A, et al. PD-L1 expression by neurons nearby tumors indicates better prognosis in glioblastoma patients. Journal of Neuroscience. 2013;33(35):14231–45.

46. Xue S, Hu M, Li P, Ma J, Xie L, Teng F, et al. Relationship between expression of PD-L1 and tumor angiogenesis, proliferation, and invasion in glioma. Oncotarget. 2017;8(30):49702–12.

47. Antonios JP, Soto H, Everson RG, Moughon D, Orpilla JR, Shin NP, et al. Immunosuppressive tumor-infltrating myeloid cells mediate adaptive immune resistance via a PD-1/PD-L1 mechanism in glioblastoma. Neuro-Oncology. 2017;19(6):796–807.

48. Berghoff AS, Kiesel B, Widhalm G, Rajky O, Ricken G, Wohrer A, et al. Programmed death ligand 1 expression and tumor-infiltrating lymphocytes in glioblastoma. Neuro-Oncology. 2015;17(8):1064–75.

49. D’Arrigo P, Russo M, Rea A, Tufano M, Guadagno E, Del Basso De Caro ML, et al. A regulatory role for the co-chaperone FKBP51s in PD-L1 expression in glioma. Oncotarget. 2017;8(40):1–21.

50. DiDomenico J, Lamano JB, Oyon D, Li Y, Veliceasa D, Kaur G, et al. The immune checkpoint protein PD-L1 induces and maintains regulatory T cells in glioblastoma. OncoImmunology. 2018;7(7).

51. Garber ST, Hashimoto Y, Weathers SP, Xiu J, Gatalica Z, Verhaak RGW, et al. Immune checkpoint blockade as a potential therapeutic target: Surveying CNS malignancies. Neuro-Oncology. 2016;18(10):1357–66.

52. Han J, Hong Y, Lee YS. PD-L1 expression and combined status of PD-L1/PD-1-positive tumor infiltrating mononuclear cell density predict prognosis in glioblastoma patients. Journal of Pathology and Translational Medicine. 2017;51(1):40–8.

53. Heiland DH, Haaker G, Delev D, Mercas B, Masalha W, Heynckes S, et al. Comprehensive analysis of PD-L1 expression in glioblastoma multiforme. Oncotarget. 2017;8(26):42214–25.

54. Heynckes S, Daka K, Franco P, Gaebelein A, Frenking JH, Doria-Medina R, et al. Crosslink between Temozolomide and PD-L1 immune-checkpoint inhibition in glioblastoma multiforme. BMC Cancer. 2019;19(1).

55. Heynckes S, Gaebelein A, Haaker G, Grauvogel J, Franco P, Mader I, et al. Expression differences of programmed death ligand 1 in de-novo and recurrent glioblastoma multiforme. Oncotarget. 2017;8(43):74170–7.

56. Lou Y, Shi J, Guo D, Qureshi AK, Song L. Function of PD-L1 in antitumor immunity of glioma cells. Saudi Journal of Biological Sciences. 2017;24(4):803–7.

57. Miyazaki T, Ishikawa E, Matsuda M, Akutsu H, Osuka S, Sakamoto N, et al. Assessment of PD-1 positive cells on initial and secondary resected tumor specimens of newly diagnosed glioblastoma and its implications on patient outcome. Journal of Neuro-Oncology. 2017;133(2):277–85.

58. Qiu XY, Hu DX, Chen WQ, Chen RQ, Qian SR, Li CY, et al. PD-L1 confers glioblastoma multiforme malignancy via Ras binding and Ras/Erk/EMT activation. Biochimica et Biophysica Acta - Molecular Basis of Disease. 2018;1864(5):1754–69.

59. Ricklefs FL, Alayo Q, Krenzlin H, Mahmoud AB, Speranza MC, Nakashima H, et al. Immune evasion mediated by PD-L1 on glioblastoma-derived extracellular vesicles. Science Advances. 2018;4(3).

60. Tamura R, Tanaka T, Ohara K, Miyake K, Morimoto Y, Yamamoto Y, et al. Persistent restoration to the immunosupportive tumor microenvironment in glioblastoma by bevacizumab. Cancer Science. 2019;110(2):499–508.

61. Wilmotte R, Burkhardt K, Kindler V, Belkouch MC, Dussex G, De Tribolet N, et al. B7-homolog I expression by human glioma: A new mechanism of immune evasion. NeuroReport. 2005;16(10):1081–5.

62. Xiu J, Piccioni D, Juarez T, Pingle SC, Hu J, Rudnick J, et al. Multi-platform molecular profiling of a large cohort of glioblastomas reveals potential therapeutic strategies. Oncotarget. 2016;7(16):21556–69.

63. De Waele J, Marcq E, Van Audenaerde JRM, Van Loenhout J, Deben C, Zwaenepoel K, et al. Poly(I:C) primes primary human glioblastoma cells for an immune response invigorated by PD-L1 blockade. OncoImmunology. 2018;7(3).

64. Takashima Y, Kawaguchi A, Kanayama T, Hayano A, Yamanaka R. Correlation between lower balance of Th2 helper T-cells and expression of PD-L1/PD-1 axis genes enables prognostic prediction in patients with glioblastoma. Oncotarget. 2018;9(27):19065–78.

65. Yao Y, Luo F, Tang C, Chen D, Qin Z, Hua W, et al. Molecular subgroups and B7-H4 expression levels predict responses to dendritic cell vaccines in glioblastoma: an exploratory randomized phase II clinical trial. Cancer Immunology, Immunotherapy. 2018;67(11):1777–88.

66. Gabrusiewicz K, Li X, Wei J, Hashimoto Y, Marisetty AL, Ott M, et al. Glioblastoma stem cell-derived exosomes induce M2 macrophages and PD-L1 expression on human monocytes. OncoImmunology. 2018;7(4).

67. Berghoff AS, Kiesel B, Widhalm G, Wilhelm D, Rajky O, Kurscheid S, et al. Correlation of immune phenotype with IDH mutation in diffuse glioma. Neuro-Oncology. 2017;19(11):1460–8.

68. Yao Y, Tao R, Wang X, Wang Y, Mao Y, Zhou LF. B7-H1 is correlated with malignancy-grade gliomas but is not expressed exclusively on tumor stem-like cells. Neuro-Oncology. 2009;11(6):757–66.

69. Hodges TR, Ott M, Xiu J, Gatalica Z, Swensen J, Zhou S, et al. Mutational burden, immune checkpoint expression, and mismatch repair in glioma: Implications for immune checkpoint immunotherapy. Neuro-Oncology. 2017;19(8):1047–57.

70. Jang BS, Kim IA. A radiosensitivity gene signature and PD-L1 predict the clinical outcomes of patients with lower grade glioma in TCGA. Radiotherapy and Oncology. 2018;128(2):245–53.

71. Samson A, Scott KJ, Taggart D, West EJ, Wilson E, Nuovo GJ, et al. Intravenous delivery of oncolytic reovirus to brain tumor patients immunologically primes for subsequent checkpoint blockade. Science Translational Medicine. 2018;10(422).

72. Zeng J, Zhang XK, Chen HD, Zhong ZH, Wu QL, Lin SX. Expression of programmed cell death-ligand 1 and its correlation with clinical outcomes in gliomas. Oncotarget. 2016;7(8):8944–55.

73. Gatalica Z, Xiu J, Swensen J, Vranic S. Molecular characterization of cancers with NTRK gene fusions. Modern Pathology. 2019;32(1):147–53.

74. Silginer M, Nagy S, Happold C, Schneider H, Weller M, Roth P. Autocrine activation of the IFN signaling pathway may promote immune escape in glioblastoma. Neuro-Oncology. 2017;19(10):1338–49.

75. Wang Y, Wang L. miR-34a attenuates glioma cells progression and chemoresistance via targeting PD-L1. Biotechnology Letters. 2017;39(10):1485–92.

76. Zhang Y, Pan C, Wang J, Cao J, Liu Y, Wang Y, et al. Genetic and immune features of resectable malignant brainstem gliomas. Oncotarget. 2017;8(47):82571–82.

77. Parsa AT, Waldron JS, Panner A, Crane CA, Parney IF, Barry JJ, et al. Loss of tumor suppressor PTEN function increases B7-H1 expression and immunoresistance in glioma. Nature Medicine. 2007;13(1):84–8.

78. Hwang K, Koh EJ, Choi EJ, Kang TH, Han JH, Choe G, et al. PD-1/PD-L1 and immune-related gene expression pattern in pediatric malignant brain tumors: clinical correlation with survival data in Korean population. Journal of Neuro-Oncology. 2018;139(2):281–91.

79. Kozlowska AK, Tseng HC, Kaur K, Topchyan P, Inagaki A, Bui VT, et al. Resistance to cytotoxicity and sustained release of interleukin-6 and interleukin-8 in the presence of decreased interferon-γ after differentiation of glioblastoma by human natural killer cells. Cancer Immunology, Immunotherapy. 2016;65(9):1085–97.

80. Speranza MC, Passaro C, Ricklefs F, Kasai K, Klein SR, Nakashima H, et al. Preclinical investigation of combined gene-mediated cytotoxic immunotherapy and immune checkpoint blockade in glioblastoma. Neuro-Oncology. 2018;20(2):225–35.

81. Han SJ, Ahn BJ, Waldron JS, Yang I, Fang S, Crane CA, et al. Gamma interferon-mediated superinduction of B7-H1 in PTEN-deficient glioblastoma: A paradoxical mechanism of immune evasion. NeuroReport. 2009;20(18):1597–602.

82. Ivanov VN, Wu J, Wang TJC, Hei TK. Inhibition of ATM kinase upregulates levels of cell death induced by cannabidiol and γ-irradiation in human glioblastoma cells. Oncotarget. 2019;10(8):825–46.

83. Song X, Shao Y, Jiang T, Ding Y, Xu B, Zheng X, et al. Radiotherapy Upregulates Programmed Death Ligand-1 through the Pathways Downstream of Epidermal Growth Factor Receptor in Glioma. EBioMedicine. 2018;28:105–13.

84. Nduom EK, Wei J, Yaghi NK, Huang N, Kong LY, Gabrusiewicz K, et al. PD-L1 expression and prognostic impact in glioblastoma. Neuro-Oncology. 2016;18(2):195–205.

85. Li S, Zhang W, Wu C, Gao H, Yu J, Wang X, et al. HOXC10 promotes proliferation and invasion and induces immunosuppressive gene expression in glioma. FEBS Journal. 2018;285(12):2278–91.

86. Chen H, Shi Z, Gao B, Fu F, Zhang X. Elevated expression of B7-H6 in U87 cells-derived glioma stem like cells is associated with biological characteristics. Xi bao yu fen zi mian yi xue za zhi = Chinese journal of cellular and molecular immunology. 2016;32(9):1168–73.

87. Wintterle S, Schreiner B, Mitsdoerffer M, Schneider D, Chen L, Meyermann R, et al. Expression of the B7-Related Molecule B7-H1 by Glioma Cells: A Potential Mechanism of Immune Paralysis. Cancer Research. 2003;63(21):7462–7.

88. Li D, Cheng S, Zou S, Zhu D, Zhu T, Wang P, et al. Immuno-PET Imaging of 89Zr Labeled Anti-PD-L1 Domain Antibody. Molecular Pharmaceutics. 2018;15(4):1674–81.

89. Xiao ZX, Chen RQ, Hu DX, Xie XQ, Yu SB, Chen XQ. Identification of repaglinide as a therapeutic drug for glioblastoma multiforme. Biochemical and Biophysical Research Communications. 2017;488(1):33–9.

90. Tran CT, Wolz P, Egensperger R, Kösel S, Imai Y, Bise K, et al. Differential expression of MHC class II molecules by microglia and neoplastic astroglia: Relevance for the escape of astrocytoma cells from immune surveillance. Neuropathology and Applied Neurobiology. 1998;24(4):293–301.

91. Anderson RC, Elder JB, Brown MD, Mandigo CE, Parsa AT, Kim PD, et al. Changes in the immunologic phenotype of human malignant glioma cells after passaging in vitro. Clinical Immunology. 2002;102(1):84–95.

92. Wu AH, Wang YJ, Zhang X, Low WC. Expression of immune-related molecules in glioblastoma multiform cells. Chinese Journal of Cancer Research. 2003;15(2):112–5.

93. Anderson RCE, Anderson DE, Elder JB, Brown MD, Mandigo CE, Parsa AT, et al. Lack of B7 expression, not human leukocyte antigen expression, facilitates immune evasion by human malignant gliomas. Neurosurgery. 2007;60(6):1129–36.

94. Pan D, Wei X, Liu M, Feng S, Tian X, Feng X, et al. Adenovirus mediated transfer of p53, GM-CSF and B7-1 suppresses growth and enhances immunogenicity of glioma cells. Neurological Research. 2010;32(5):502–9.

95. Parney IF, Waldron JS, Parsa AT. Flow cytometry and in vitro analysis of human glioma-associated macrophages: Laboratory investigation. Journal of Neurosurgery. 2009;110(3):572–82.

96. Xie T, Liu B, Dai CG, Lu ZH, Dong J, Huang Q. Glioma stem cells reconstruct similar immunoinflammatory microenvironment in different transplant sites and induce malignant transformation of tumor microenvironment cells. Journal of Cancer Research and Clinical Oncology. 2019;145(2):321–8.

97. Ulasov IV, Rivera AA, Han Y, Curiel DT, Zhu ZB, Lesniak MS. Targeting adenovirus to CD80 and CD86 receptors increases gene transfer efficiency to malignant glioma cells. Journal of Neurosurgery. 2007;107(3):617–27.

98. Joki T, Kikuchi T, Akasaki Y, Saitoh S, Abe T, Ohno T. Induction of effective antitumor immunity in a mouse brain tumor model using B7-1 (CD80) and intercellular adhesive molecule 1 (ICAM-1; CD54) transfection and recombinant interleukin 12. International Journal of Cancer. 1999;82(5):714–20.

99. Mu YG, Peng H, Zhang JY, Shao CJ, Wu CY, Chen ZP. Expression of costimulator 4-1BBL and B7-1 on glioma cell lines. Ai zheng = Aizheng = Chinese journal of cancer. 2006;25(3):326–9.

100. Naganuma H, Sasaki A, Satoh E, Nagasaka M, Nakano S, Isoe S, et al. Down-regulation of transforming growth factor-β and interleukin-10 secretion from malignant glioma cells by cytokines and anticancer drugs. Journal of Neuro-Oncology. 1998;39(3):227–36.

101. Li G, Pan W, Yang X, Miao J. Gene co-expression network and function modules in three types of glioma. Molecular Medicine Reports. 2015;11(4):3055–63.

102. Parney IF, Farr-Jones MA, Chang LJ, Petruk KC. Human glioma immunobiology in vitro: Implications for immunogene therapy. Neurosurgery. 2000;46(5):1169–78.

103. Parney IF, Farr-Jones MA, Koshal A, Chang LJ, Petruk KC, Liu CY, et al. Human brain tumor cell culture characterization after immunostimulatory gene transfer. Neurosurgery. 2002;50(5):1094–102.

104. Parney IF, Petruk KC, Zhang C, Farr-Jones M, Sykes DB, Chang LJ. Granulocyte-macrophage colony-stimulating factor and B7-2 combination immunogene therapy in an allogeneic Hu-PBL-SCID/beige mouse-human glioblastoma multiforme model. Human Gene Therapy. 1997;8(9):1073–85.

105. Schreiner B, Wischhusen J, Mitsdoerffer M, Schneider D, Bornemann A, Melms A, et al. Expression of the B7-Related Molecule ICOSL by Human Glioma Cells In Vitro and In Vivo. GLIA. 2003;44(3):296–301.

106. Zhou Z, Luther N, Ibrahim GM, Hawkins C, Vibhakar R, Handler MH, et al. B7-H3, a potential therapeutic target, is expressed in diffuse intrinsic pontine glioma. Journal of Neuro-Oncology. 2013;111(3):257–64.

107. Li RG, Gao Z, Jiang Y. B7-H3 repression by miR-539 suppresses cell proliferation in human gliomas. International Journal of Clinical and Experimental Pathology. 2017;10(4):4363–9.

108. Liu YT. Expressions and clinical significance of PDCD4 and B7-H4 in optic gliomas in children. International Eye Science. 2014;14(8):1391–3.

109. Yao Y, Ye H, Qi Z, Mo L, Yue Q, Baral A, et al. B7–H4(B7x)-mediated cross-talk between glioma-initiating cells and macrophages via the IL6/JAK/STAT3 pathway lead to poor prognosis in glioma patients. Clinical Cancer Research. 2016;22(11):2778–90.

110. Yao Y, Wang X, Jin K, Zhu J, Wang Y, Xiong S, et al. B7–H4 is preferentially expressed in non-dividing brain tumor cells and in a subset of brain tumor stem-like cells. Journal of Neuro-Oncology. 2008;89(2):121–9.

111. Mo LJ, Ye HX, Mao Y, Yao Y, Zhang JM. B7-H4 expression is elevated in human U251 glioma stem-like cells and is inducible in monocytes cultured with U251 stem-like cell conditioned medium. Chinese Journal of Cancer. 2013;32(12):653–60.

112. Kaneko S, Nakatani Y, Takezaki T, Hide T, Yamashita D, Ohtsu N, et al. Ceacam1L modulates STAT3 signaling to control the proliferation of glioblastoma-initiating cells. Cancer Research. 2015;75(19):4224–34.

113. Li J, Liu X, Duan Y, Wang H, Su W, Wang Y, et al. Abnormal expression of circulating and tumor-infiltrating carcinoembryonic antigen-related cell adhesion molecule 1 in patients with glioma. Oncology Letters. 2018;15(3):3496–503.

114. Castriconi R, Daga A, Dondero A, Zona G, Poliani PL, Melotti A, et al. NK cells recognize and kill human glioblastoma cells with stem cell-like properties. Journal of Immunology. 2009;182(6):3530–9.

115. Chandramohan V, Bryant JD, Piao H, Keir ST, Lipp ES, Lefaivre M, et al. Validation of an immunohistochemistry assay for detection of CD155, the poliovirus receptor, in malignant gliomas. Archives of Pathology and Laboratory Medicine. 2017;141(12):1697–704.

116. Gromeier M, Lachmann S, Rosenfeld MR, Gutin PH, Wimmer E. Intergeneric poliovirus recombinants for the treatment of malignant glioma. Proceedings of the National Academy of Sciences of the United States of America. 2000;97(12):6803–8.

117. Merrill MK, Bernhardt G, Sampson JH, Wikstrand CJ, Bigner DD, Gromeier M. Poliovirus receptor CD155-targeted oncolysis of glioma. Neuro-Oncology. 2004;6(3):208–17.

118. Maherally Z, Smith JR, An Q, Pilkington GJ. Receptors for hyaluronic acid and poliovirus: A combinatorial role in glioma invasion? PLoS ONE. 2012;7(2).

119. Sloan KE, Eustace BK, Stewart JK, Zehetmeier C, Torella C, Simeone M, et al. CD155/PVR plays a key role in cell motility during tumor cell invasion and migration. BMC Cancer. 2004;4.

120. Yang X, Chen E, Jiang H, Muszynski K, Harris RD, Giardina SL, et al. Evaluation of IRES-mediated, cell-type-specific cytotoxicity of poliovirus using a colorimetric cell proliferation assay. Journal of Virological Methods. 2009;155(1):44–54.

121. Enloe BM, Jay DG. Inhibition of Necl-5 (CD155/PVR) reduces glioblastoma dispersal and decreases MMP-2 expression and activity. Journal of Neuro-Oncology. 2011;102(2):225–35.

