## supplementary figures for "Expression of inhibitory checkpoint ligands by Glioblastoma Multiforme cells and the implications of an enhanced stem cell-like phenotype"

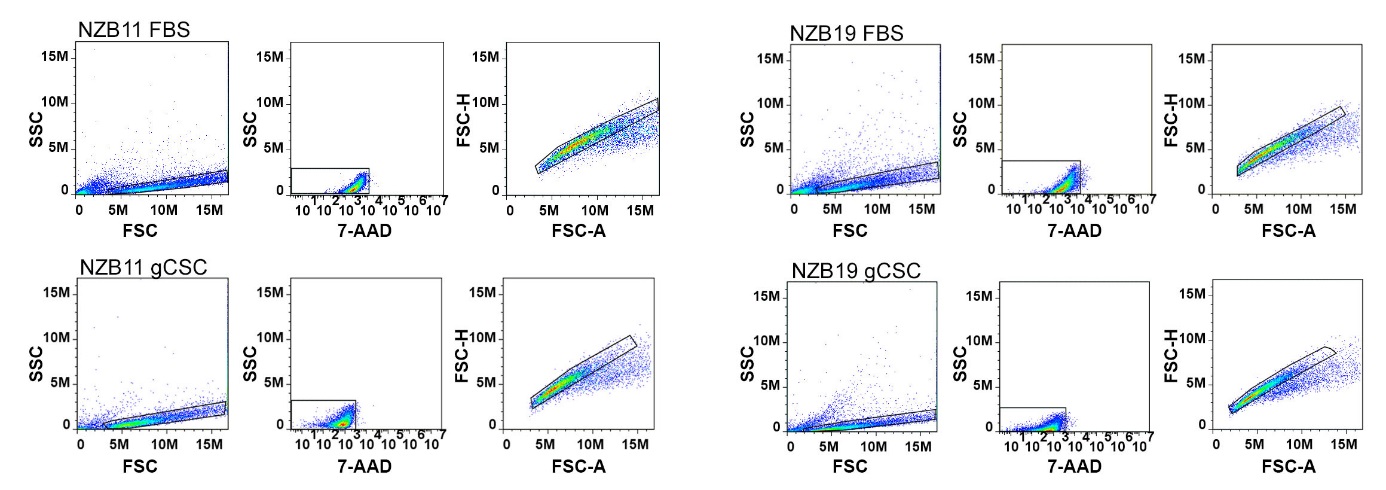

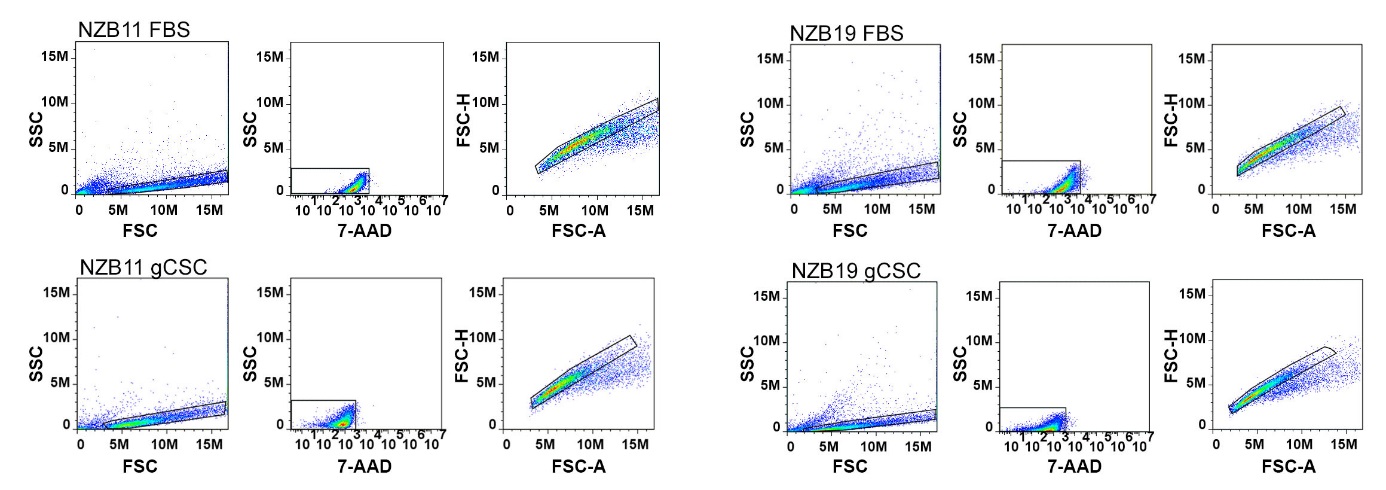

Figure S1 Gating of respective primary New Zealand Glioblastoma cell lines for flow cytometry analysis. NZB11 and NZB19 serum-cultured and gCSC cells were gated using forward vs. side-scatter to delineate the cell population. 7-AAD was used to gate out dead cells as per a right-ward shift in the population above that of auto-fluorescence. Doublets were gated out as per forward scatter height vs. area.

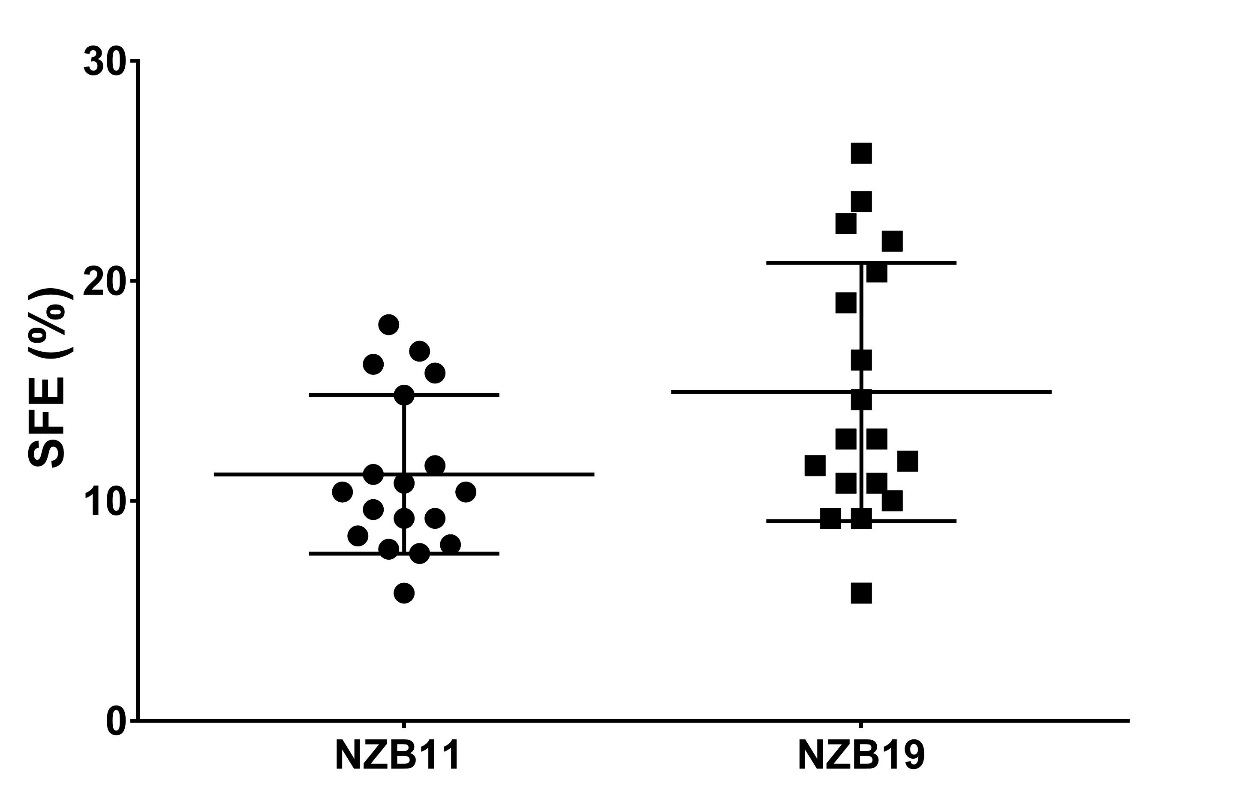

Figure S2 Sphere forming efficiency. NZB11 and NZB19 glioma-spheres were serially passaged for three generations. Generations two and three were seeded at limited dilutions of 5 cells/µL. Spheres were cultured for 14 days and then counted to determine sphere forming efficiencies (SFE). Shown are three independent repeats.

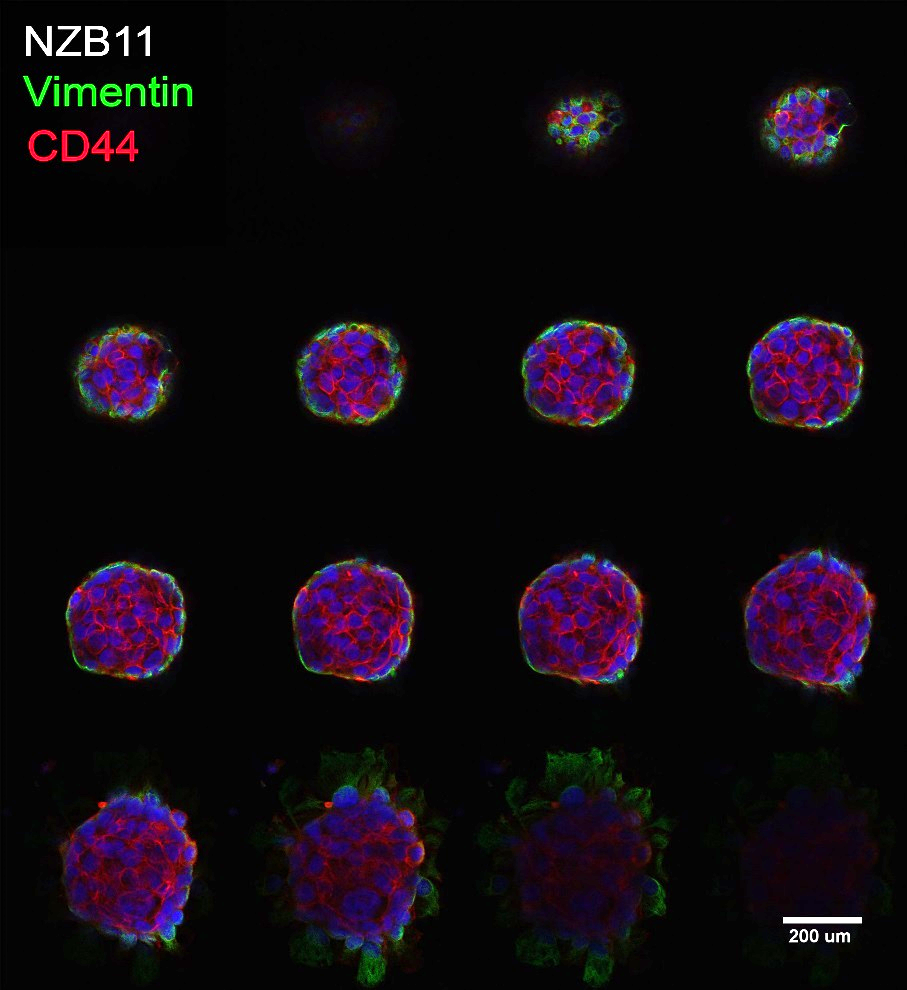

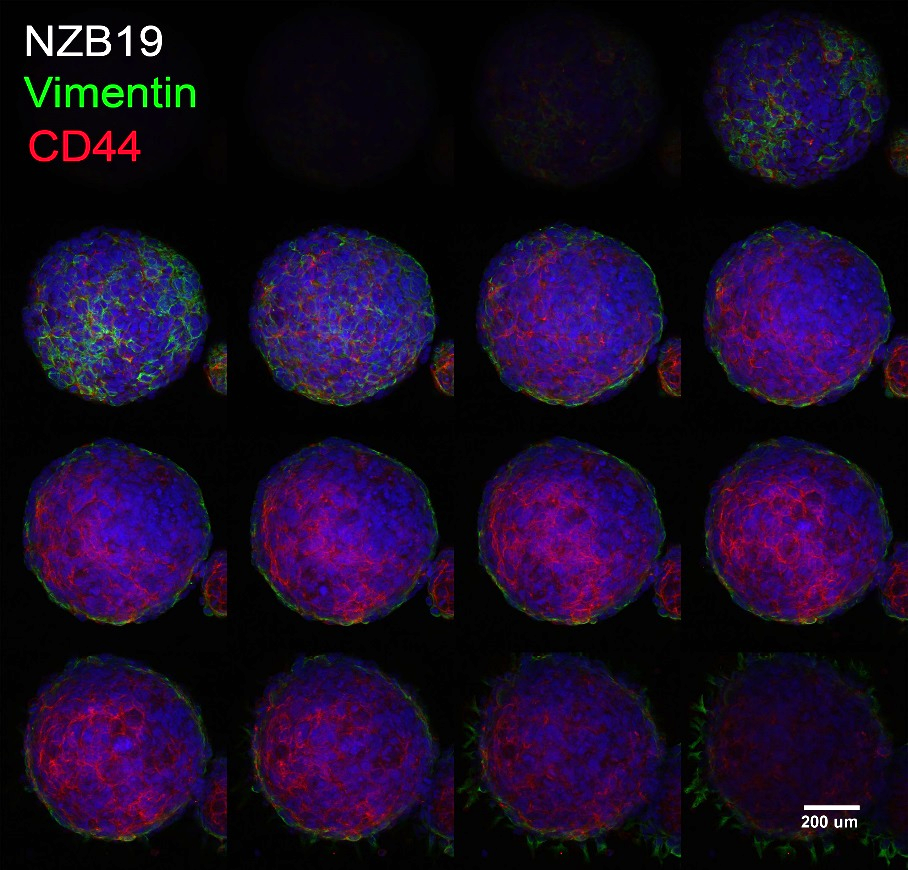

Figure S3 Confocal Z-stack montages of CD44 and Vimentin expression in glioma-spheres. NZB11 and NZB19 glioma-sphere localisation of CD44 and Vimentin expression within gCSC derived glioma-spheres. NZB11 slice thickness = 0.47µm. NZB19 slice thickness = 1.18µm. Images were acquired at x40 magnification. Scale bar = 200µm.

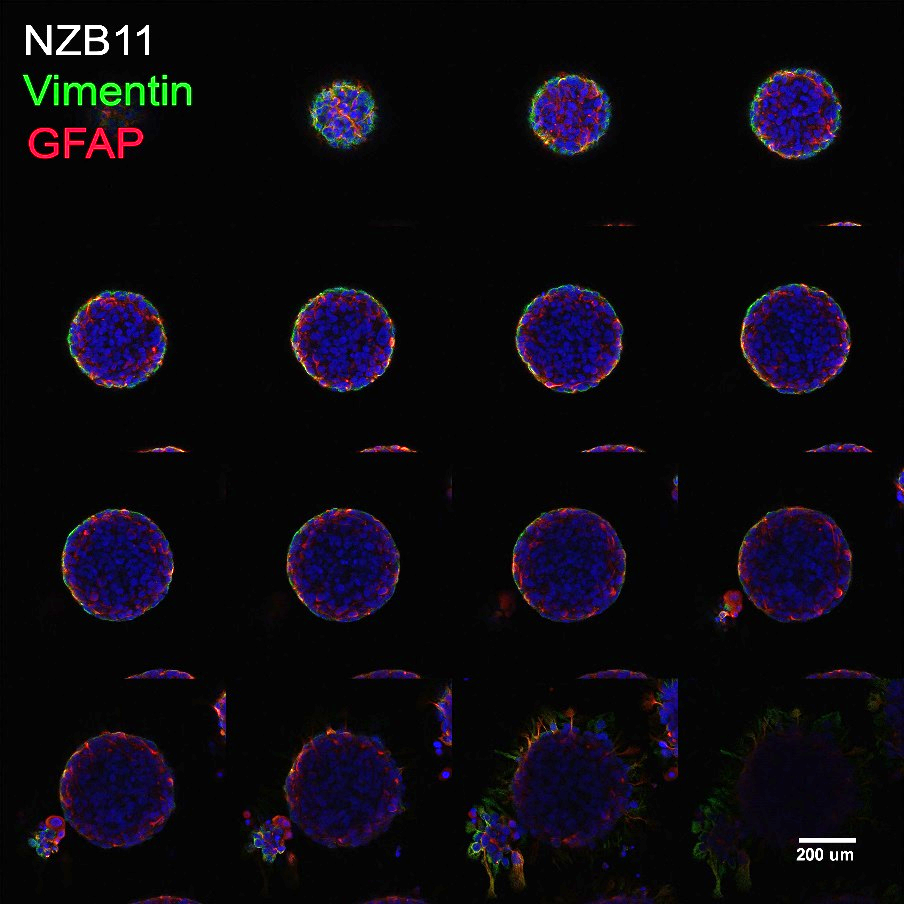

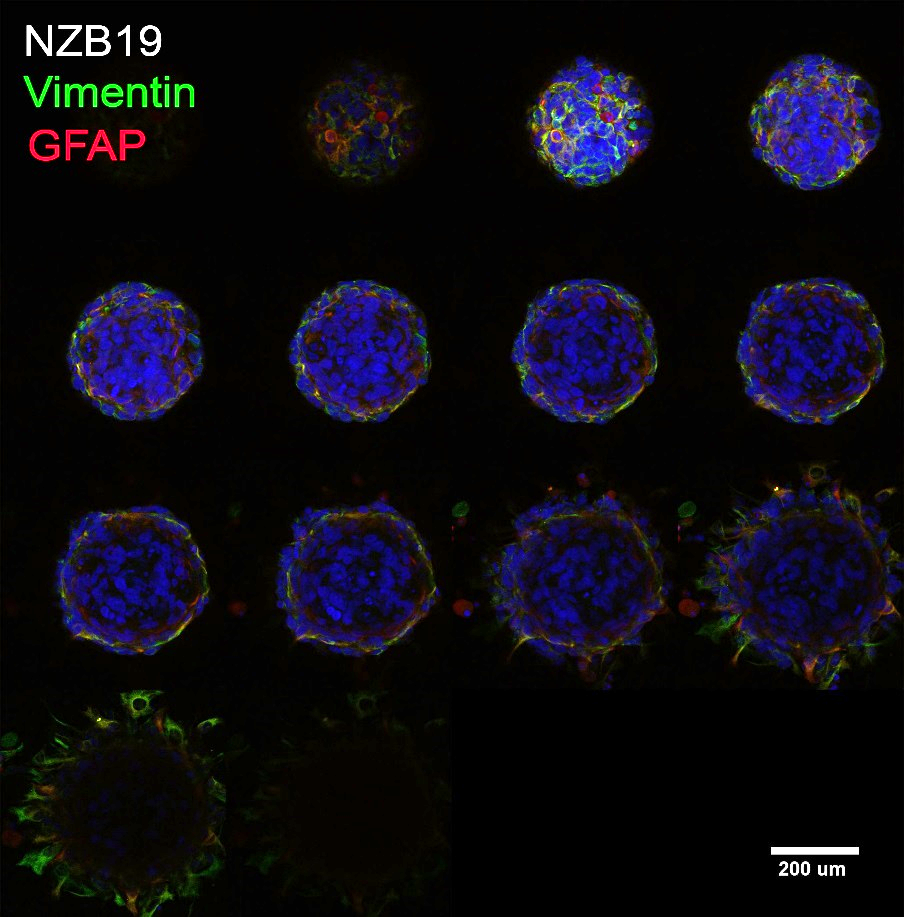

Figure S4 Confocal Z-stack montages of GFAP and Vimentin expression in glioma-spheres. NZB11 and NZB19 glioma-sphere localisation of GFAP and Vimentin expression within gCSC derived glioma-spheres. NZB11 slice thickness = 1.10µm. NZB19 slice thickness = 1.09µm. Images were acquired at x40 magnification. Scale bar = 200µm.

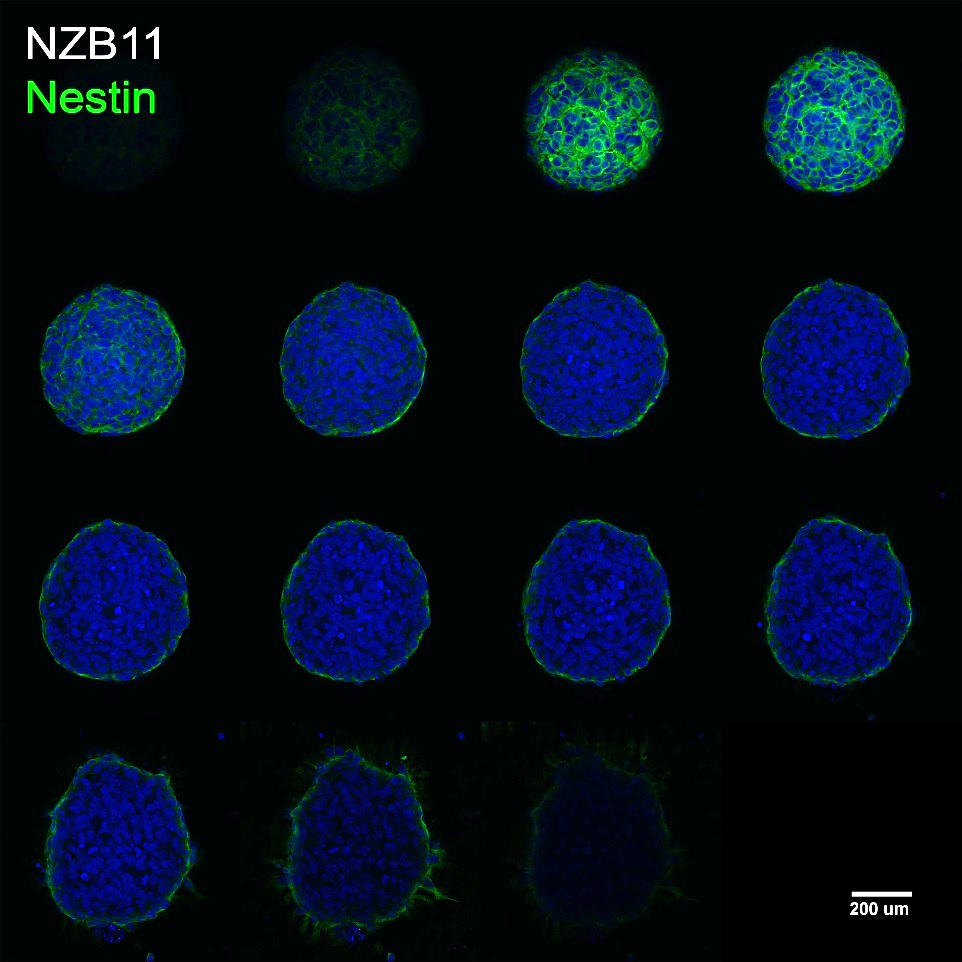

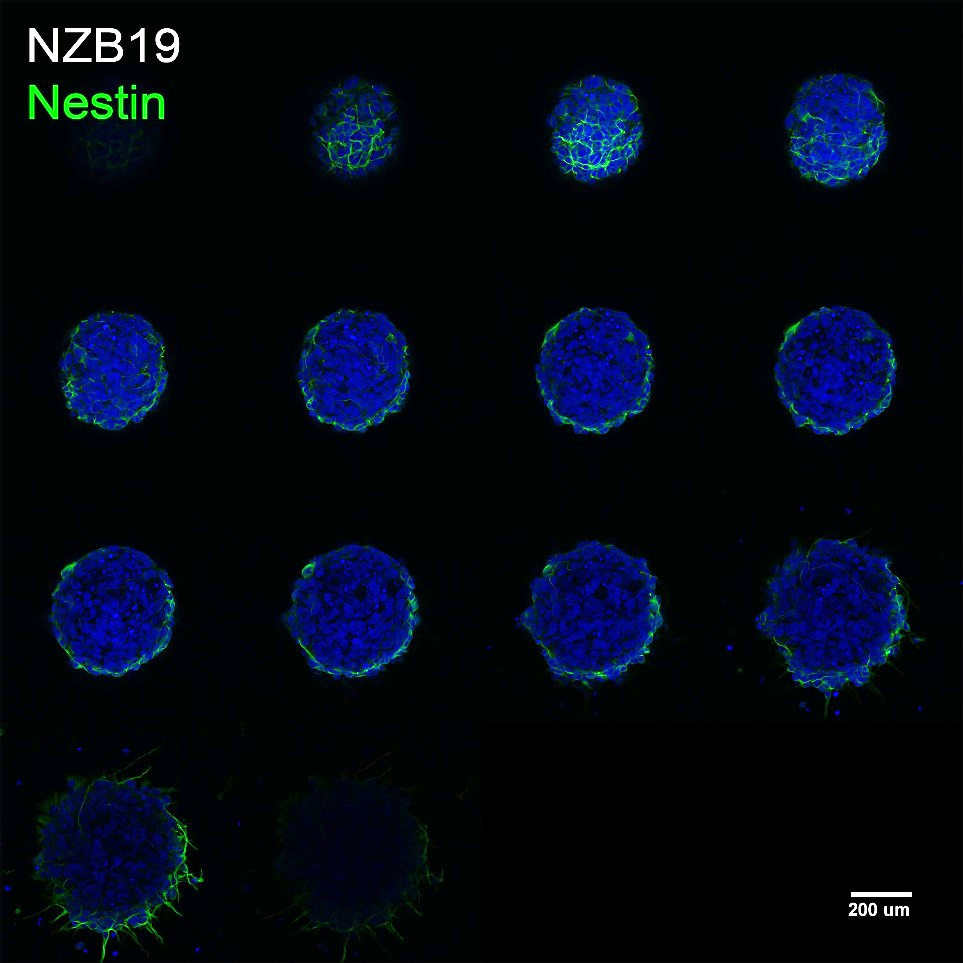

Figure S5 Confocal Z-stack montages of Nestin expression in glioma-spheres. NZB11 and NZB19 glioma-sphere localisation of Nestin expression within gCSC derived glioma-spheres. NZB11 slice thickness = 1.19µm. NZB19 slice thickness = 1.09µm. Images were acquired at x40 magnification. Scale bar = 200µm.

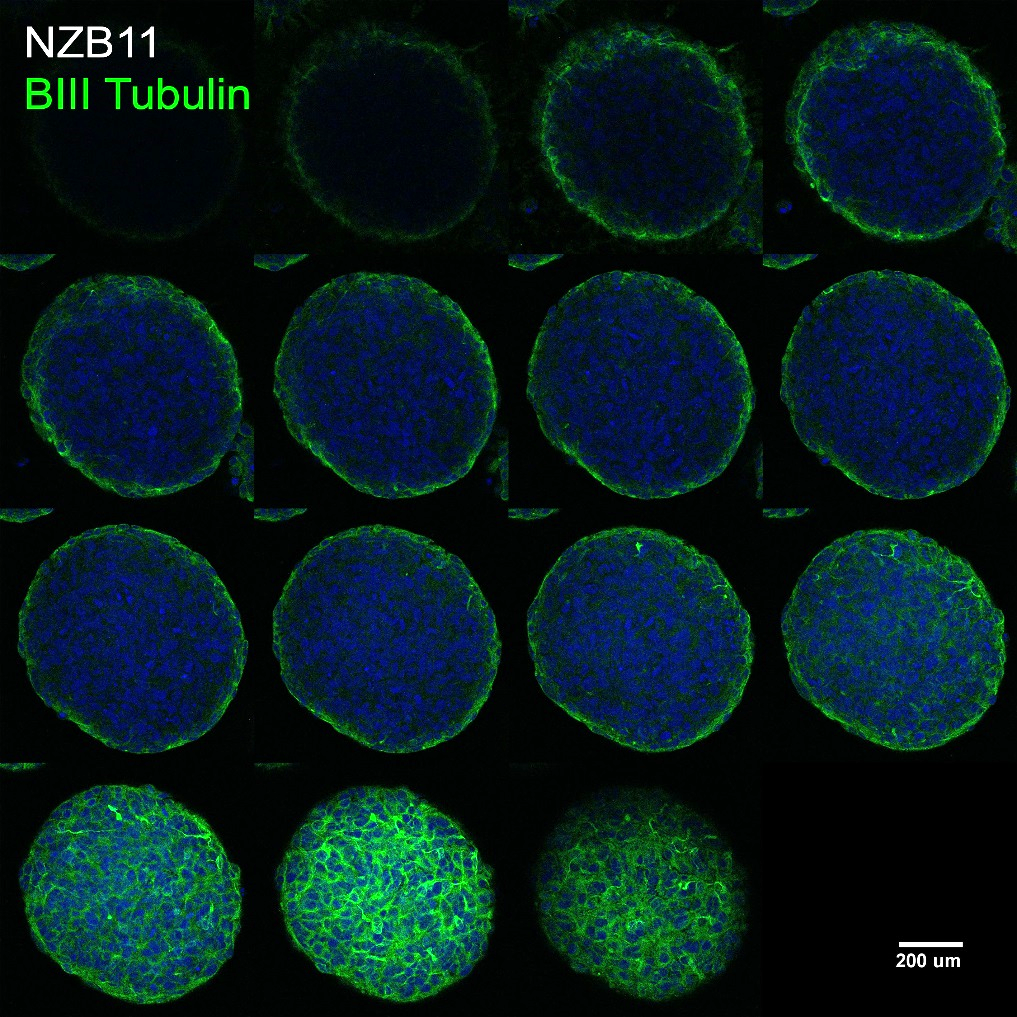

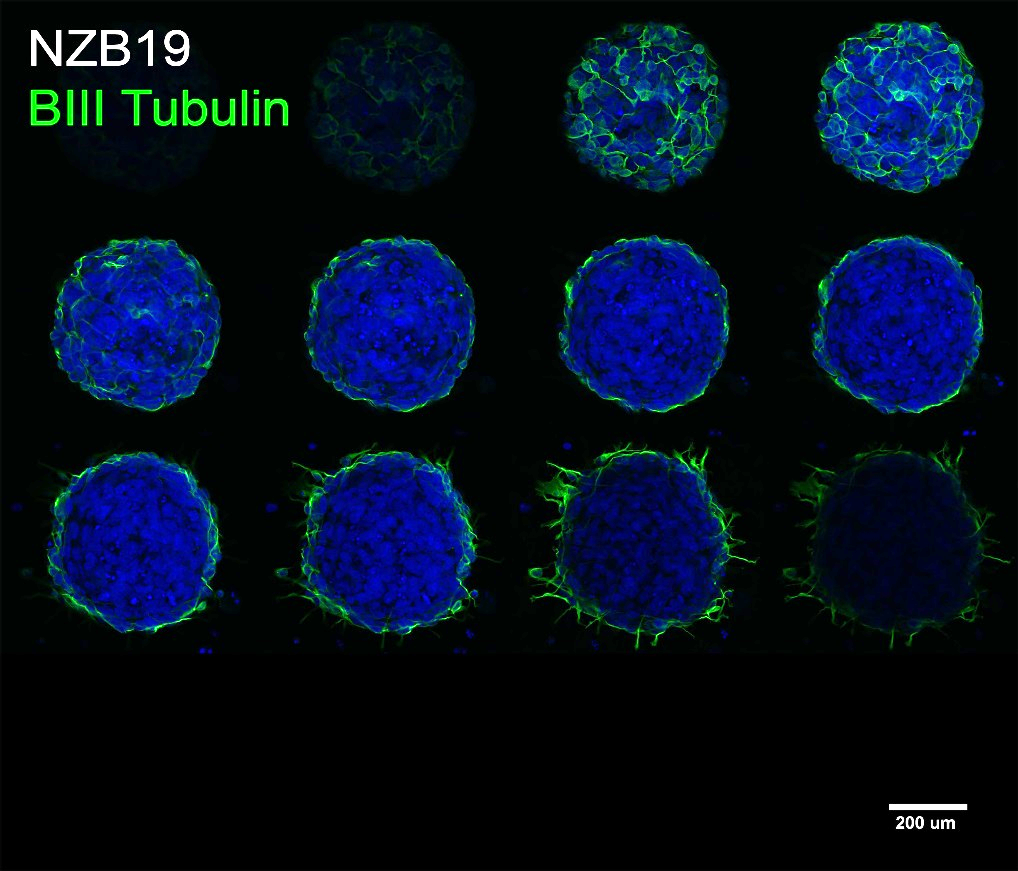

Figure S6 Confocal Z-stack montages of BIII Tubulin expression in glioma-spheres. NZB11 and NZB19 glioma-sphere localisation of BIII Tubulin Vimentin expression within gCSC derived glioma-spheres. NZB11 slice thickness = 1.23µm. NZB19 slice thickness = 1.06µm. Images were acquired at x40 magnification. Scale bar = 200µm.

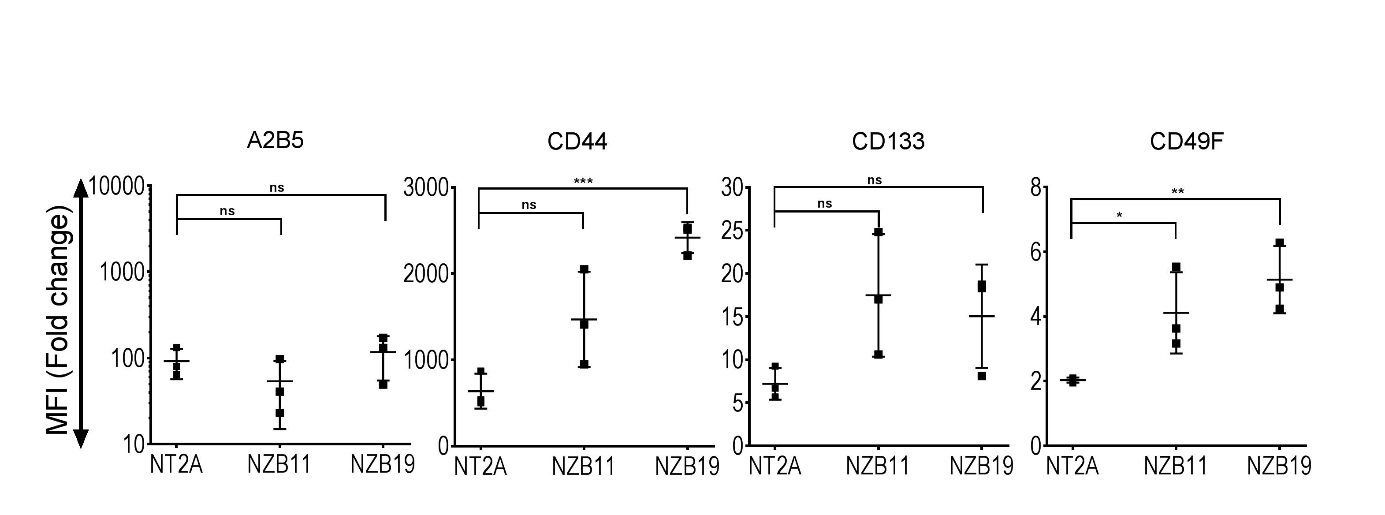

Figure S7 Stem cell associated marker expression analysis across two primary New Zealand GBM cell lines. Median fluorescent intensity fold change from auto-fluorescence for each glioma stem cell marker. Side by side comparisons of NZB11 and NZB19 serum-cultured cells vs. NT2A astrocytic control. Shown are three independent repeats. Paired students T-test analysis was carried out and respective P values are shown. P = 0.05 (*), 0.01 (**), 0.001 (***), 0.0001(****).

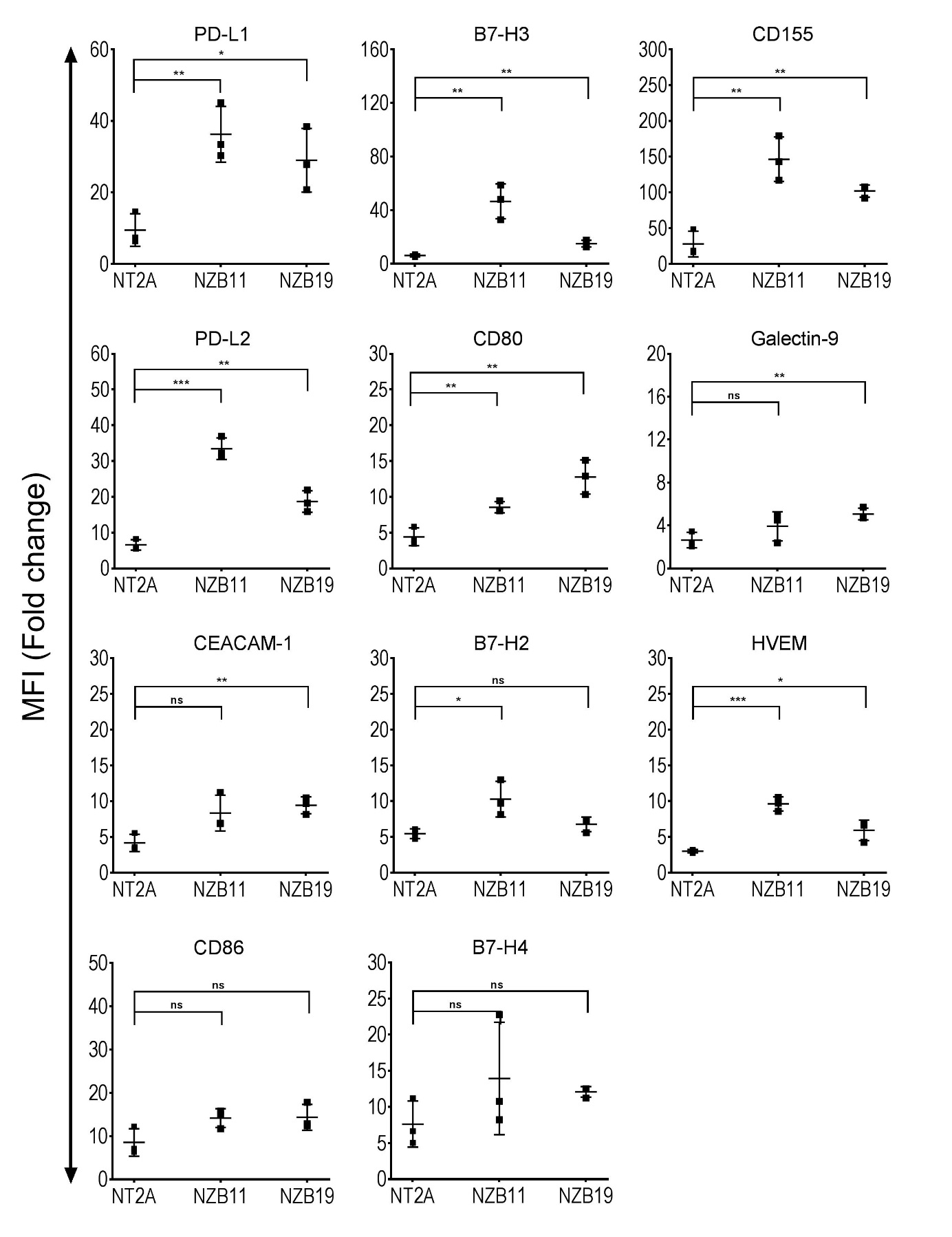

Figure S8 Checkpoint ligand expression analysis across two primary New Zealand GBM cell lines. Median fluorescent intensity fold change from auto-fluorescence for each checkpoint ligand. Side by side comparisons of serum-cultured NZB11 and NZB19 cells vs. NT2A astrocytic control. Shown are three independent repeats. Paired students T-test analysis was carried out and respective P values are shown. P = 0.05 (*), 0.01 (**), 0.001 (***), 0.0001(****).

Table S1 Reported expression of checkpoint ligands in human glioma

| **PD-L1** |  |  |  |
| --- | --- | --- | --- |
| **Tissue** |  |  |  |
| *Classification* | *Expressed* | *Technique* | *Reference* |
| GBM**** | Yes | IHC, TMA, WB, qPCR**, RNA-seq, in-situ | (11, 22, 35, 43-65) |
| GBM**** | No | IHC | (66) |
| Astrocytoma* | Yes | IHC, WB | (46, 61, 67, 68) |
| Glioma*** | Yes | IHC, WB, qPCR**, RNA-seq, TMA, IP | (22, 35, 47, 49, 51, 56, 69-77) |
| Paediatric malignant brain tumour | Yes | IHC, NS | (78) |
| **Cells** |  |  |  |
| *Classification* | *Expressed* | *Technique* | *Reference* |
| Primary | Yes | qPCR, FC, WB, LRA, ICC | (54, 55, 61, 63, 68, 71, 75, 77) |
| Glioma cancer stem | Yes | FC, WB, ICC, qPCR | (59, 79, 80) |
| U87 | Yes | ICC, FC, qPCR, WB | (74, 77, 81-87) |
| T98G | Yes | ICC, FC, qPCR, WB | (74, 82, 87) |
| SF767 | Yes | FC, qPCR, WB | (77, 81) |
| U118 | Yes | ICC, FC, qPCR, WB | (82) |
| LN229 | Yes | IHC*****, FC, qPCR | (56, 87-89) |
| SF126 | Yes | qPCR, FC, WB | (77) |
| SF188 | Yes | qPCR, FC, WB | (77) |
| SF210 | Yes | qPCR, FC, WB | (77) |
| U251 | Yes | qPCR, FC, WB | (49, 77, 83, 85, 87) |
| U373 | Yes | qPCR, FC, WB | (77, 87) |
| D54 | Yes | ICC, FC, qPCR, WB | (49) |
| LN308 | Yes | FC, qPCR | (56) |
| LN18 | Yes | FC, qPCR | (87) |
| U138 | Yes | FC, qPCR | (87) |
| LN428 | Yes | FC, qPCR | (87) |
| LN319 | Yes | FC, qPCR | (87) |
| D247 | Yes | FC, qPCR | (87) |

| **PD-L2** |  |  |  |
| --- | --- | --- | --- |
| **Tissue** |  |  |  |
| *Classification* | *Expressed* | *Technique* | *Reference* |
| GBM**** | Yes | TMA, IHC, RNA-seq | (22, 34, 35, 63, 64) |
| Glioma*** | Yes | RNA-seq, IHC | (22, 34) |
| **Cells** |  |  |  |
| *Classification* | *Expressed* | *Technique* | *Reference* |
| Primary | Yes | FC, qPCR | (63) |
| Glioma cancer stem | Yes | FC | (79) |
| U87 | Yes | ICC, FC, qPCR, WB | (82, 85) |
| T98G | Yes | ICC, FC, qPCR, WB | (82) |
| SF188 | Yes | ICC, FC, qPCR, WB | (82) |
| U251 | Yes | WB | (85) |

| **CD80** |  |  |  |
| --- | --- | --- | --- |
| **Tissue** |  |  |  |
| *Classification* | *Expressed* | *Technique* | *Reference* |
| GBM**** | Yes | TMA, RNA-seq | (22, 35) |
| GBM**** | No | IHC | (87, 90) |
| Glioma*** | Yes | TMA, RNA-seq | (22, 35) |
| Glioma*** | No | IHC | (87, 90) |
| **Cells** |  |  |  |
| *Classification* | *Expressed* | *Technique* | *Reference* |
| Primary | Yes | FC | (91) |
| Primary | No | FC | (92-95) |
| Glioma cancer stem | No | WB | (96) |
| U87 | Yes | FC | (97) |
| U87 | No | FC, qPCR | (87, 92, 94, 98) |
| T98G | No | FC, qPCR | (87, 99, 100) |
| SF767 | No | FC | (99) |
| U118 | No | FC | (98) |
| LN229 | No | FC, qPCR | (87) |
| U251 | No | FC, qPCR | (87, 94) |
| U373 | Yes | FC | (97) |
| U373 | No | FC, qPCR | (87, 98) |
| LN308 | No | FC, qPCR | (87) |
| MGR2 | No | FC | (99) |
| MGR1 | No | FC | (99) |
| SKMG4 | No | FC | (99) |
| UW28 | No | FC | (99) |
| A172 | No | FC, qPCR | (87, 100) |
| KG-1-C | No | FC | (100) |
| YMG2 | No | FC | (100) |
| YMG4 | No | FC | (100) |
| LN18 | No | FC, qPCR | (87) |
| U138 | No | FC, qPCR | (87) |
| LN428 | No | FC, qPCR | (87) |
| LN319 | No | FC, qPCR | (87) |
| D247 | No | FC, qPCR | (87) |

| **CD86** |  |  |  |
| --- | --- | --- | --- |
| **Tissue** |  |  |  |
| *Classification* | *Expressed* | *Technique* | *Reference* |
| GBM**** | Yes | TMA | (101) |
| Glioma*** | No | IHC | (87) |
| **Cells** |  |  |  |
| *Classification* | *Expressed* | *Technique* | *Reference* |
| Primary | No | FC | (91-93, 95, 102) |
| U87 | Yes | FC | (97) |
| U87 | No | FC, qPCR | (87, 92) |
| T98G | No | FC, qPCR | (87) |
| D54 | No | FC | (103, 104) |
| LN229 | No | FC, qPCR | (87) |
| U251 | No | FC, qPCR | (87) |
| U373 | Yes | FC | (97) |
| U373 | No | FC, qPCR | (87) |
| LN308 | No | FC, qPCR | (87) |
| A172 | No | FC, qPCR | (87) |
| LN18 | No | FC, qPCR | (87) |
| U138 | No | FC, qPCR | (87) |
| LN428 | No | FC, qPCR | (87) |
| LN319 | No | FC, qPCR | (87) |
| D247 | No | FC, qPCR | (87) |

| **B7-H2** |  |  |  |
| --- | --- | --- | --- |
| **Tissue** |  |  |  |
| *Classification* | *Expressed* | *Technique* | *Reference* |
| GBM**** | Yes | IHC | (105) |
| Astrocytoma* | Yes | IHC | (105) |
| **Cells** |  |  |  |
| *Classification* | *Expressed* | *Technique* | *Reference* |
| Primary | Yes | FC, qPCR | (105) |

| **B7-H3** |  |  |  |
| --- | --- | --- | --- |
| **Tissue** |  |  |  |
| *Classification* | *Expressed* | *Technique* | *Reference* |
| GBM**** | Yes | IHC | (42) |
| Glioma*** | Yes | qPCR, IHC, TMA | (33, 41, 42, 106) |
| Astrocytoma* | Yes | IHC, PCR | (33) |
| **Cells** |  |  |  |
| *Classification* | *Expressed* | *Technique* | *Reference* |
| Glioma cancer stem | Yes | N/A | (86) |
| U87 | Yes | qPCR, WB | (107) |
| LN229 | Yes | FC, qPCR, WB | (33) |
| U251 | Yes | qPCR, WB | (107) |

| **B7-H4** |  |  |  |
| --- | --- | --- | --- |
| **Tissue** |  |  |  |
| *Classification* | *Expressed* | *Technique* | *Reference* |
| GBM**** | Yes | IHC | (65) |
| Paediatric optic glioma | Yes | IHC | (108) |
| **Cells** |  |  |  |
| *Classification* | *Expressed* | *Technique* | *Reference* |
| Primary | Yes | ELISA, FC, qPCR | (109) |
| Glioma cancer stem | Yes | FC, ICC | (86, 110) |
| U251 | Yes | ICC, FC | (111) |

| **CEACAM-1** |  |  |  |
| --- | --- | --- | --- |
| **Tissue** |  |  |  |
| *Classification* | *Expressed* | *Technique* | *Reference* |
| GBM**** | Yes | IHC | (112) |
| Glioma*** | Yes | IHC | (113) |
| **Cells** |  |  |  |
| *Classification* | *Expressed* | *Technique* | *Reference* |
| Primary | Yes | ICC, qPCR, WB | (112) |

| **CD155** |  |  |  |
| --- | --- | --- | --- |
| **Tissue** |  |  |  |
| *Classification* | *Expressed* | *Technique* | *Reference* |
| GBM**** | Yes | IHC, WB | (114, 115) |
| Glioma*** | Yes | IHC | (116, 117) |
| **Cells** |  |  |  |
| *Classification* | *Expressed* | *Technique* | *Reference* |
| Primary | Yes | ICC, FC, WB | (117, 118) |
| Glioma cancer stem | Yes | ICC, FC | (114) |
| U87 | Yes | ICC, WB, ELISA, FC | (119, 120) |
| A172 | Yes | ICC, WB | (121) |
| U251 | Yes | ICC, WB | (121) |

*Pilocytic, diffuse and anaplastic astrocytoma, ** qPCR = qPCR, qRT-PCR, Taqman, *** Grade I, II, III. Brainstem glioma, **** GBM, Grade IV glioma, ***** Mouse transplant
